# The ancestral population size conditioned on the reconstructed phylogenetic tree with occurrence data

**DOI:** 10.1101/755561

**Authors:** Marc Manceau, Ankit Gupta, Timothy Vaughan, Tanja Stadler

**Affiliations:** Department of Biosystems Science and Engineering, ETH Zürich, Basel, Switzerland

**Keywords:** birth-death process, fossilized birth-death model, epidemiology, macroevolution, phylogenetics

## Abstract

We consider a homogeneous birth-death process with three different sampling schemes. First, individuals can be sampled through time and included in a reconstructed tree. Second, they can be sampled through time and only recorded as a point ‘occurrence’ along a timeline. Third, extant individuals are sampled and included in the reconstructed tree with a fixed probability. We further consider that sampled individuals can be removed or not from the process, upon sampling, with fixed probability.

Given an outcome of the process, composed of the joint observation of a reconstructed phylogenetic tree and a record of occurrences not included in the tree, we derive the conditional probability distribution of the population size any time in the past. We additionally provide an algorithm to simulate ancestral population size trajectories given the observation of a reconstructed tree and occurrences.

This distribution can readily be used to perform inferences of the ancestral population size in the field of epidemiology and macroevolution. In epidemiology, these results will pave the way towards jointly considering data from case count studies and reconstructed transmission trees. In macroevolution, it will foster the joint examination of the fossil record and extant taxa to reconstruct past biodiversity.

## 1. Introduction

Since seminal papers by Yule (1925), Kendall (1948), and much later by Nee et al. (1994), birth-death models have become ubiquitous in evolutionary biology. They are used as an underlying tree prior, or as an underlying population dynamics prior in recent studies spanning fields as diverse as macroevolution, linguistics, or epidemiology (see e.g. Heath et al., 2014; Gray et al., 2009; Ratmann et al., 2016). Computing reliable estimates of the ancestral number of species, languages or infected individuals, i.e. estimates of past population size, is key to the understanding of past processes in these fields.

In both macroevolution and epidemiology, these inferences have initially relied on the fossil record and the case counts record, modeled as a sampling of individuals from the full process through time (Foote, 2000; Starrfelt and Liow, 2016).

In the recent years, impressive sequencing efforts targeting present-day species and pathogens have enabled the reconstruction of phylogenies, the branch-lengths of which can be used to infer ancestral population dynamics. Two main approaches have been introduced to estimate past population sizes from reconstructed evolutionary trees.

The first one, in a maximum likelihood framework, consists of estimating the birth and death rate of the population using all branch-lengths of the tree ahead. Analytical estimates of past population size are then computed in a second step, based on simplifications of the data, such as, e.g, taking only into account the number of individuals at the beginning (Morlon et al., 2011) or more recently at the beginning and end of the process (Billaud et al., 2019).

The second one is a variation of the first method in a Bayesian setting. Birth and death rates are supposed to be piecewise constant with a given *a priori* distribution, and their posterior is sampled using all branch-lengths. Past population sizes are then computed by integrating the expected number of individuals over the rates posterior distribution (Stadler et al., 2013).

While this second method accounts for uncertainty in the estimated population sizes, it also relies on numerical Monte-Carlo methods which make estimates much more computationally expensive to compute. Additionally, the statistical signal in the number of lineages through time is only used to compute rate estimates, whereas it could in principle also more precisely inform the past population size distribution.

Statistical approaches stemming from the analysis of case count data or from the analysis of reconstructed evolutionary trees have been part of separate literature bodies for years, historically yielding conflict between biodiversity estimates based on the fossil record and estimates based on phylogenies of extant taxa (Quental and Marshall, 2010 but see also Morlon et al., 2011).

A first path towards merging these disparate data was introduced by the fossilized birth-death model of Stadler (2010), who considered a birth-death model with sampling and inclusion of individuals in the tree through time. This allowed taking into account infection trees reconstructed from pathogen sequences sampled throughout an epidemic (Stadler et al., 2011). In macroevolution, it paved the way to more precise phylogenetic dating using well-conserved fossil taxa which could be placed on a reconstructed phylogeny using morphological characters (Gavryushkina et al., 2016). Not so well-conserved fossils (i.e. occurrences) have also been used with this model, using a Markov Chain Monte Carlo (MCMC) scheme to integrate over all possible placements along a fixed tree (Heath et al., 2014).

Very recently, Vaughan et al. (2019) introduced another extension to the fossilized birth-death model, considering that individuals sampled through time could either be included or not included in the reconstructed evolutionary tree. This variation has profound consequences for data analysis, for it allows one to combine the occurrence record that cannot be placed along a tree (e.g. poorly preserved fossils, or case count epidemiological record) with samples for which molecular sequences or phenotypic measurements allow reconstructing a phylogenetic tree. They also provide a Monte-Carlo particle filtering algorithm to reconstruct past population sizes by sampling from the distribution of the number of ancestors conditioned on the joint observation of the reconstructed tree and the occurrence record. Analytical developments around this new model have further been made by Gupta et al. (2019), who derived an analytical formula for the probability density of an outcome of the process.

In this paper, we build on the analytical developments presented by Gupta et al. (2019), and aim at providing a more analytical, and faster, way to compute the same distribution as originally targeted by Vaughan et al. (2019). The efficiency of our method paves the way towards considering much bigger datasets, or towards extending the method to multi-type or density-dependent birth-death processes.

In section 2, we present the model, notation, and an overview of the strategy to express the targeted distribution. In section 3, we adapt the main results of Gupta et al. (2019) to compute the probability density of observations made after a given time, conditioned on the past population size. In section 4, we provide a way to compute the joint density of the past population size and observations made before a given time. Combining results of sections 3 and 4, we compute the distribution of past population sizes conditional on the full outcome of the process in section 5. We finally discuss applications and potential extensions of the model.

## 2. Model and notation

### 2.1. Parameters of the process

We consider a population of individuals, any of which can give birth to another individual at rate *λ* or die at rate *µ*. The process starts at time *t*_*or*_ in the past with one individual, and unfolds until reaching present time 0, i.e. time is oriented from the present towards the past.

We superimpose to this background population dynamics three different sampling schemes. First, individuals can be *ψ-sampled* at rate *ψ* throughout their lifetime. When *ψ*-sampled, the individual will be included in the reconstructed tree. Second, individuals can be *ω-sampled* at rate *ω* throughout their life-time. When *ω*-sampled, the individual is not included in the reconstructed tree, but its sampling time is nevertheless recorded and thereafter called ‘an occurrence’. Last, the process is finished upon reaching the present time 0, and each extant individual at that time is *ρ-sampled* with fixed probability *ρ*, leading to their inclusion in the reconstructed tree. The sum of all per-capita rates will be called for short *γ* = *λ* + *µ* + *ψ* + *ω*.

Following Vaughan et al. (2019), we also include in the model an effect of the *ψ*- and *ω*-sampling through time on the population dynamics. We consider that, upon sampling, an individual is either removed from the process with probability *r* ∈ (0, 1), or is unaffected by the sampling with probability (1 − *r*). The overall number of individuals, denoted (*I*_*t*_), thus follows a linear birth-death process with birth rate *λ* and death rate *µ* + (*ψ* + *ω*)*r*. Because this work aims at fitting the model to empirical data, we only consider here a supercritical process with *λ > µ* + (*ψ* + *ω*)*r*.

### 2.2. Introducing useful probabilities

This process has already been thoroughly studied. In the following sections, we will use two key probabilities which analytical solutions are well known. First, we will call *u*_*t*_ the probability that a process starting at time *t* with only one individual is never sampled before reaching present time 0. We recall that *u*_*t*_ satisfies the Master equation

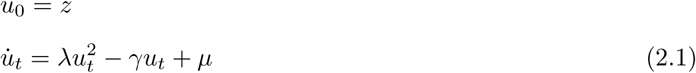

The solution of this for a particular initial condition *z* being the following

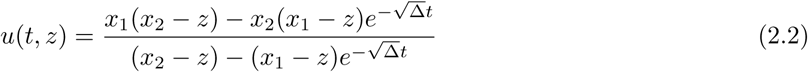

where Δ = *γ*^2^ − 4*λµ >* (*λ* + *µ*)^2^ − 4*λµ ≥* (*λ* − *µ*)^2^ *>* 0 and *x*_1_, *x*_2_ are the two roots of the polynomial *λx*^2^ − *γx* + *µ*,

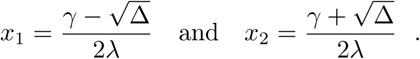

Second, we call *p*_*t*_ the probability that a process starting at time *t* with one individual leads to one sampled particle at present time 0. Writing the Master equation governing the evolution of this quantity leads to

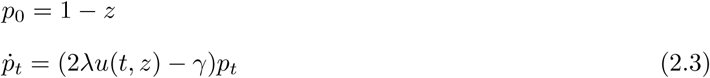

The solution of this being the following

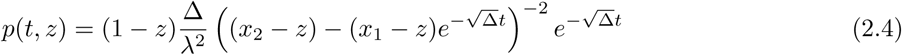

These formulas are well known, and correspond respectively to quantities called *p*_0_(*t*) and *p*_1_(*t*) in Stadler (2010). When *z* = 1 − *ρ*, we will drop the dependence on *z* and use the shorter notation *u*_*t*_, *p*_*t*_. We recall standard ways to derive these expressions in Appendix A.

### 2.3. Strategy of the paper

The process with sampling leads to the observation of two distinct objects (*𝒯*, *𝒪*), as illustrated in Figure 1.

**Figure 1:**
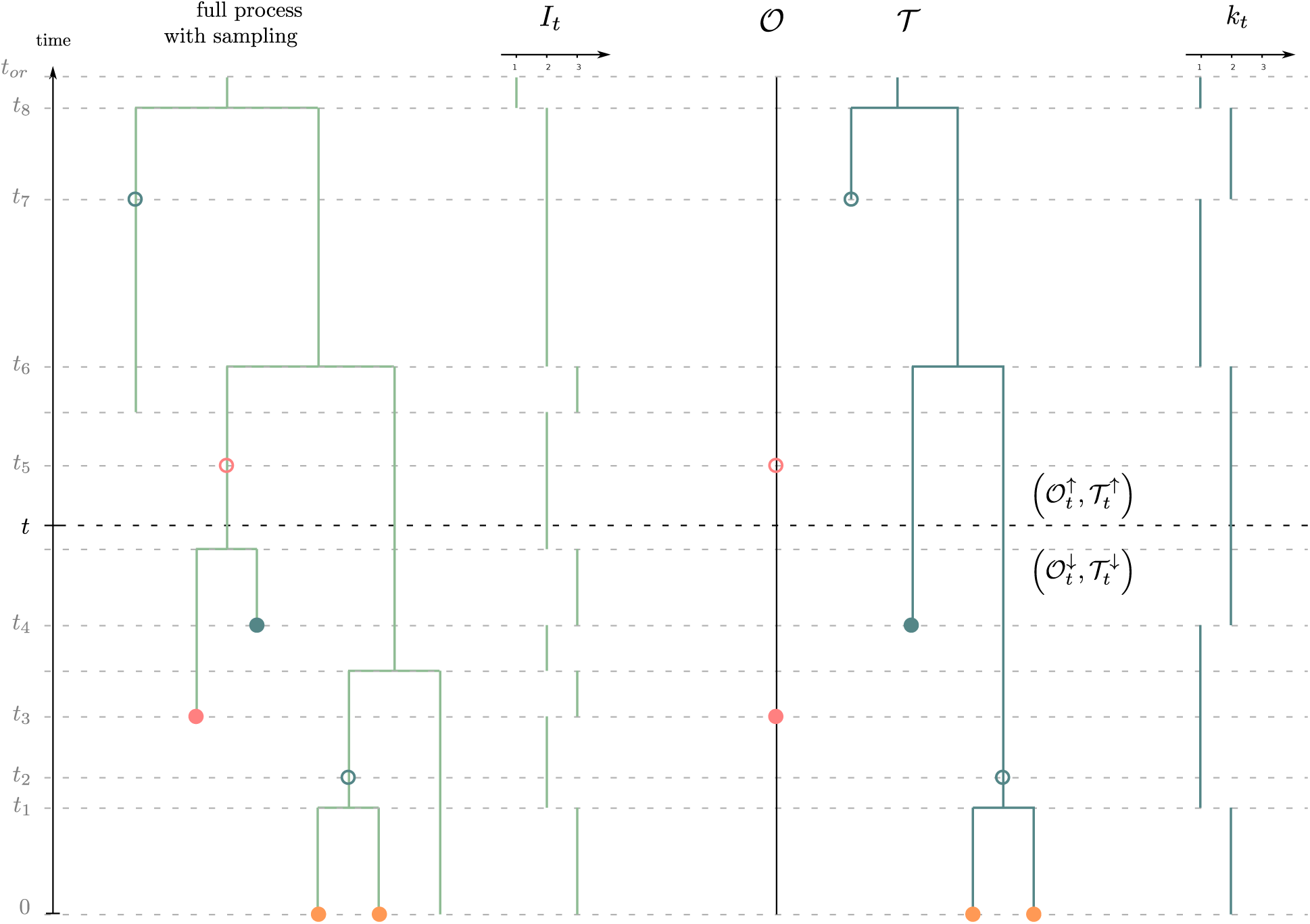
General setting of the method. To the left in green, the full process with sampling. Red dots correspond to *ω*-sampling (sampling through time without sequencing), blue dots correspond to *ψ*-sampling (sampling through time with sequencing) and orange dots correspond to *ρ*-sampling at present. Filled or unfilled dots correspond respectively to sampling with or without removal. To the right, the observations: a reconstructed tree *T* together with the record of occurrences *𝒪*.

The reconstructed tree 𝒯, on the one hand, represents the evolutionary relationships between all *ψ*-sampled and *ρ*-sampled individuals. We further consider that *ψ*-sampled individuals are labeled either as ‘removed’ or ‘non-removed’. All *ψ*-sampled removed individuals are necessarily leaves of 𝒯, whereas *ψ*-sampled non-removed ones can either stand as leaves (when the descent of the individual is not sampled) or as vertices along a branch (when the descent of the individual is further sampled), in which case they are referred to as *sampled ancestors*.

The record of occurrences *𝒪*, on the other hand, is an ordered list of all *ω*-sampling times. We also consider that these sampling times are labeled as either ‘removed’ or ‘non-removed’.

In this paper, we are interested in computing the probability distribution of the number of individuals in the past, conditioned on the observed outcome (*𝒯*, *𝒪*) of the process. If *k*_*t*_ denotes the number of sampled lineages in *𝒯* at time *t*, we call our target distribution,

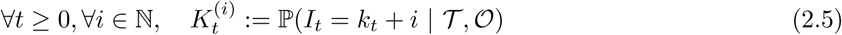

We will refer to *epochs* as the maximal time slices within which no sampling event in *𝒪*, nor branching event in *𝒯*, happened. These epochs are delimited by the union of sampling times in *𝒪*, branching times in the tree *𝒯*, and sampling times of leaves and sampled ancestors in *𝒯*. All pooled together, we call these ordered times 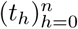, starting at present time *t*_0_ = 0 and ending at the root time *t*_*n*_ = *t*_*or*_.

At any time *t ≥* 0 we also introduce:

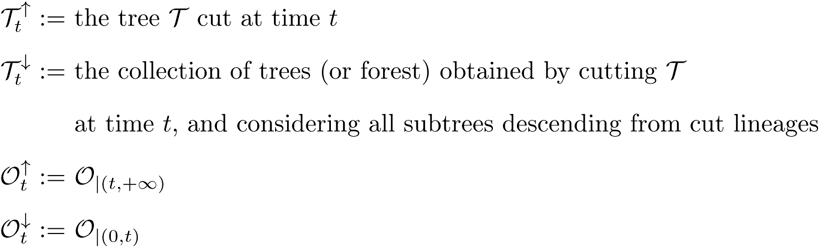

The general strategy – and outline – of the paper is the following. In a *breadth-first backward traversal* we will compute the probability density of observations made down any time *t*, conditioned on the population size at that time, as time increases towards the past. We call this probability density,

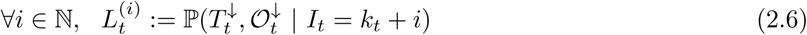

In a *breadth-first forward traversal* we will then compute the joint probability density of the observations made up to any time *t* and the population size at that time, as time decreases towards present. We call this density,

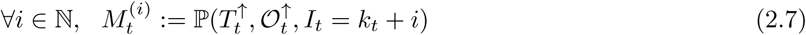

Provided we get expressions of 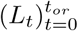 and 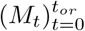, our target distribution can then be expressed by combining both, noting that:

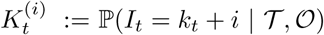

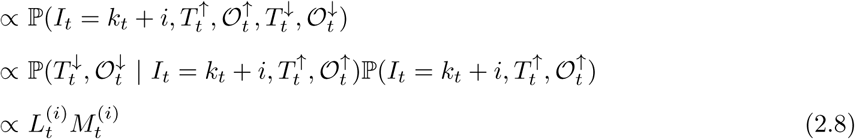

where the last line holds because, conditionally on *I*_*t*_ = *k*_*t*_ + *i*, the future of the (Markov) process is independent of what happened before.

In the process of getting the likelihood of observations under the same model, Gupta et al. (2019) provided an analytical formula and an algorithm to compute the first ingredient *L*_*t*_ in the case where all individuals are removed upon sampling (i.e. *r* = 1). We thus recall their main result, and adapt it to our slightly different framework, in the next section.

## 3. The density of observations below conditioned on past population size

We start this section by presenting the Master equations satisfied by the probability density *L*_*t*_. This provides us with a numerical algorithm to compute *L*_*t*_, which we subsequently simplify with analytical results for specific sets of parameters.

### 3.1. Master equation driving L_*t*_

One way to get the probability density *L*_*t*_ consists in studying its evolution through time, relying on its Master equation. First, remark that we can express *L*_0_ at present time 0. Indeed, provided we know the exact number of individuals living at time 0, the probability to see the tips of the tree is directly driven by the *ρ*-sampling,

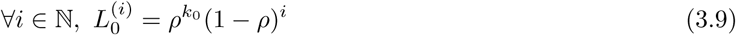

We now derive the Master equation driving the evolution of *L*_*t*_ through time across any given epoch. We consider an infinitesimal time step *δt* and list the events which could have happened in the full process between *t* and *t* + *δt*, leading to our observations. Suppose the number of observed lineages in this epoch is *k*, and the total number of individuals alive is *k* + *i*. We emphasize three cases, illustrated in Figure 2:

**Figure 2:**
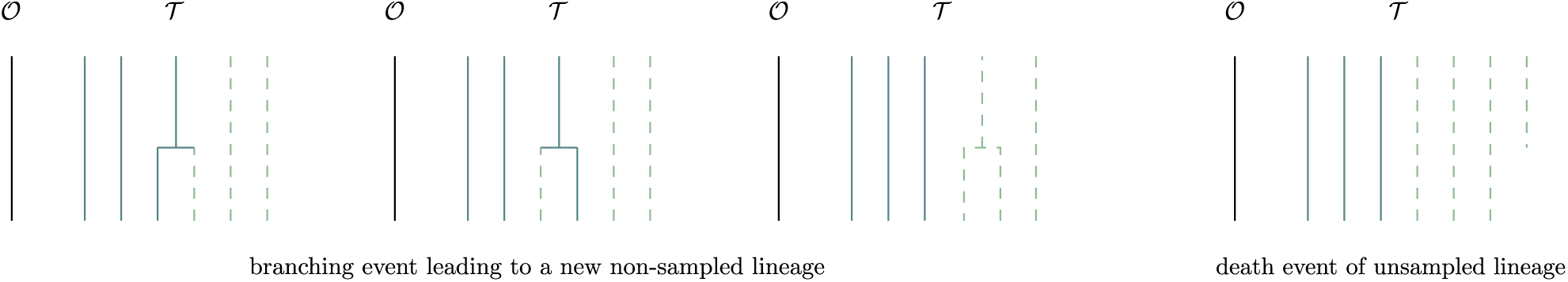
Four unobservable scenarios taken into account to derive the Master equations 3.10 and 4.25.

1. nothing happened with probability (1 − *γ*(*k* + *i*)*δt*)
2. a birth event happened
  a. among the *k* sampled lineages in 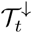, and it leads to an extinct or unsampled subtree to the left or to the right, with probability 2*λδt*.
  b. among the *i* other individuals, with probability *λiδt*.
3. a death event happened among the *i* particles, with probability *µiδt*.

These allow us to write, *∀i* ∈ ℕ,

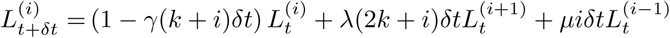

Note that for 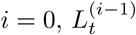 is not defined, but the term cancels out thanks to the factor *i*.

Letting *δt* → 0 and combining this with the initial condition on *L*_0_, we get the following set of ordinary differential equations driving the evolution of *L*_*t*_,

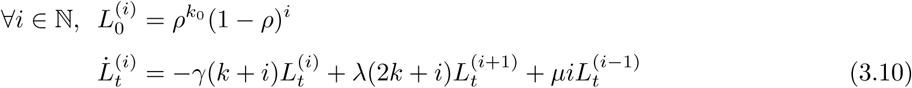

Last, we need to study how *L*_*t*_ changes at punctual events. There are 6 types of punctual events that we can come across at time *t* in the past, listed below and illustrated in Figure 3.

**Figure 3:**
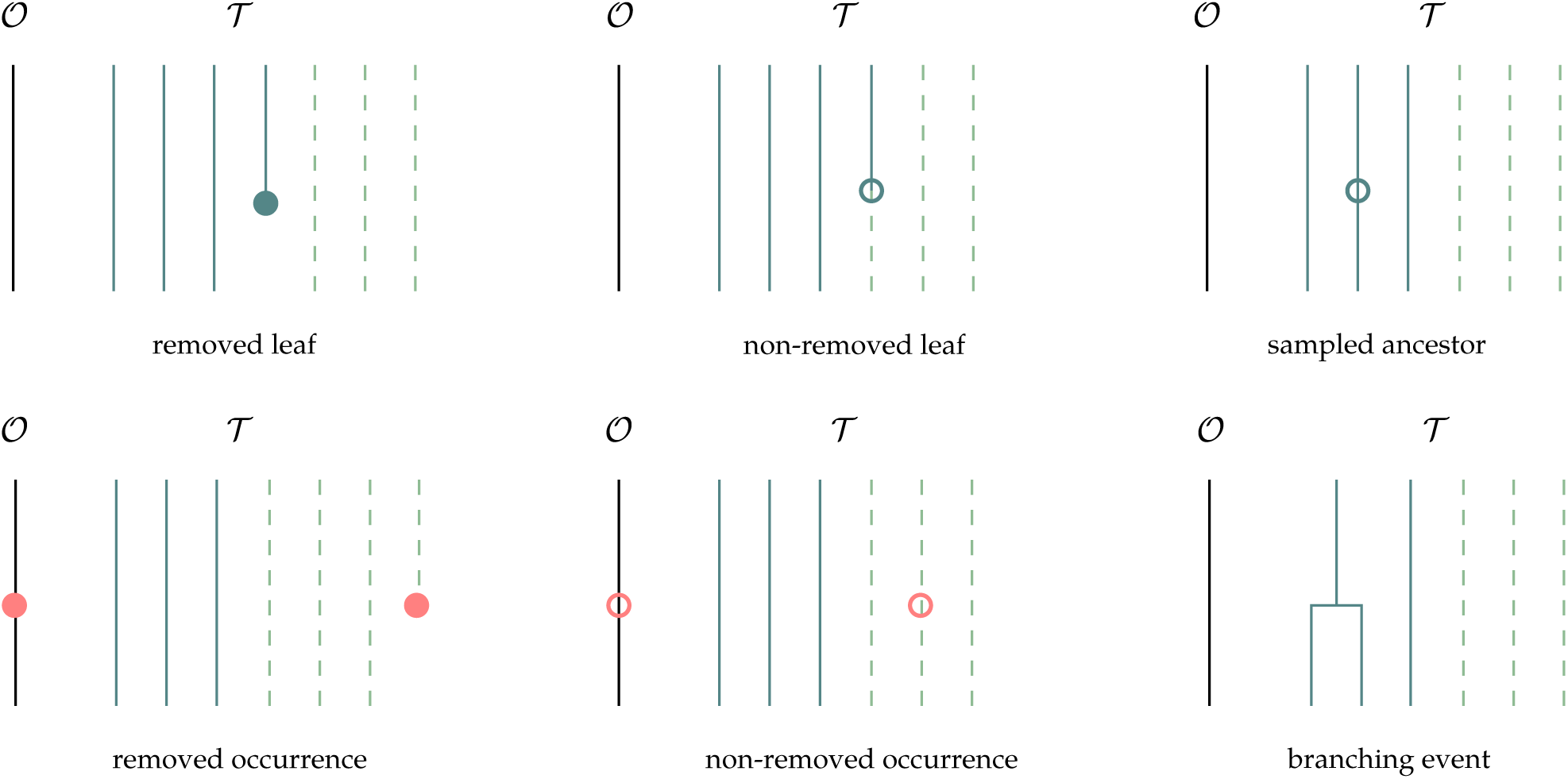
Six observable punctual events in the data.

We denote 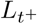 the probability just before (i.e. up) the punctual event and 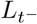 the probability immediately after (i.e. down). We can either find,

1. a leaf of 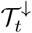, labeled as removed. This is a *ψ*-sampling with removal event for which the number of unsampled lineages remains constant, and the number of sampled lineages increases by one. It thus gives,

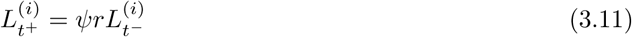
2. a leaf of 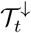, labeled as non-removed. This is a *ψ*-sampling without removal event for which one of the unsampled lineage becomes a sampled one. It thus gives,

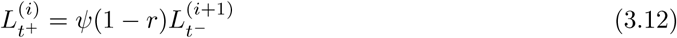
3. a sampled ancestor along a branch of 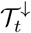, necessarily labeled as non-removed. This is a *ψ*-sampling without removal event, not impacting the number of sampled or unsampled lineages. It thus gives,

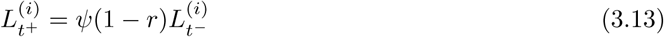
4. an occurrence in 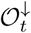, labeled as removed. This is a *ω*-sampling with removal event, for which the number of unsampled lineages increases by one. It thus gives,

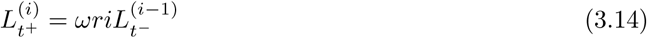 Note that here also, for 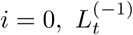 is not defined but the term cancels out thanks to the factor *i*.
5. an occurrence in 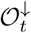, labeled as non-removed. This is a *ω*-sampling without removal event, not impacting the number of sampled or unsampled lineages. It thus gives,

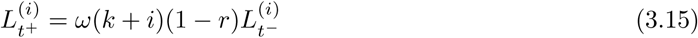
6. a branching event between two branches of 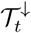. The number of sampled lineages decreases by one. It thus gives,

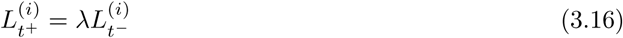

This Master equation can be numerically approximated. To do so, we fix a finite upper bound *N* on the number of hidden individuals and numerically integrate a truncated ODE system. We detail this in the following algorithm to compute an approximation of *L*_*t*_ at any time *t*.

#### Algorithm 1 Computes a numerical approximation of *L*_*t*_ for a specific set of times

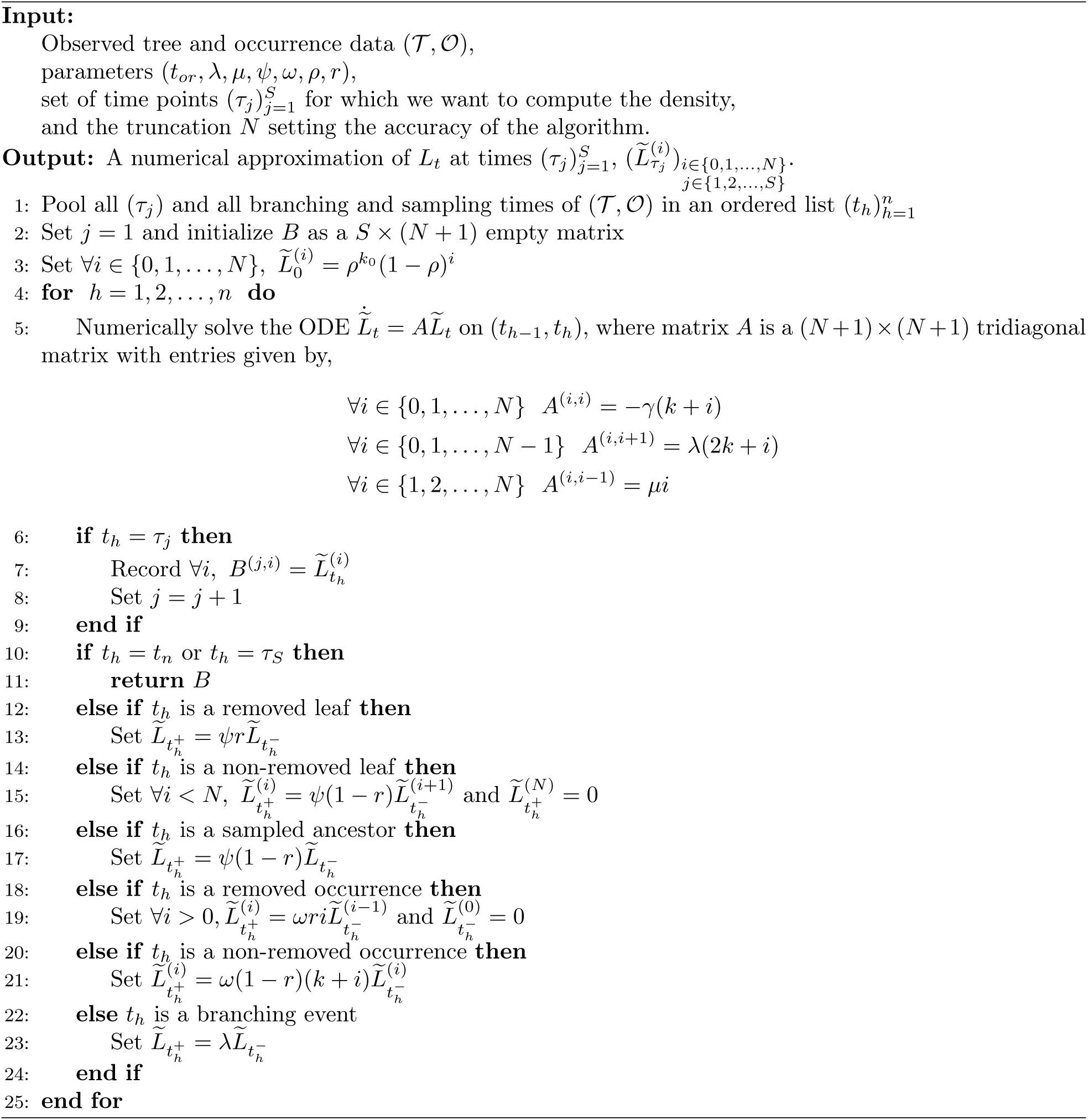

We also define a slight variation of this algorithm, that we will refer to as algorithm 1’, where no set of time points (*τ*_*j*_) is required, and the values of 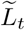 are not recorded through time (i.e. matrix *B* disappears). Instead, when reaching *t*_*n*_ = *t*_*or*_ we simply return 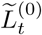, which by definition is an estimate of the probability density of (*𝒯*, *𝒪*). Note that this strategy is very close to what has been used to compute the probability density of a reconstructed tree under a logistic birth-death process (Leventhal et al., 2013).

These two algorithms will prove useful to deal with the general case. However, we can further get analytical expressions for *L*_*t*_ when *ω* = 0 as well as when *r* = 1 (Gupta et al., 2019). We expose these in the next two subsections.

### 3.2. Special case ω = 0

Suppose we can express 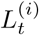 as the product 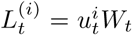 where *W*_*t*_ is a function of time only, and *u*_*t*_ is defined as in equation 2.2. We first get, from the initialization in equation (3.10), that 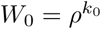. Moreover, substituting 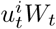 in the ODE leads to

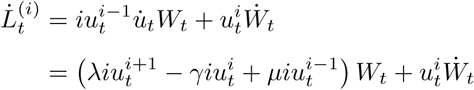

Thus leading to the following ODE for *W*_*t*_, on any epoch (*t*_*h*_, *t*_*h*+1_) where the number of sampled lineages remains fixed and equal to *k*,

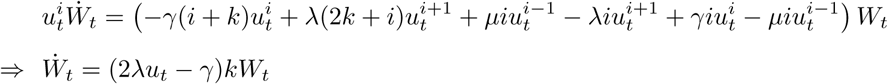

This is very close to the Master equation (2.3) governing the evolution of *p*_*t*_, and it leads to (see derivation in Appendix A),

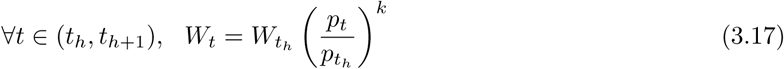

Last, because *ω* = 0, updates (3.11) to (3.16) simplify to only the following *ψ* - and *λ*-events,

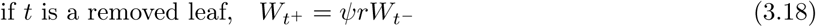

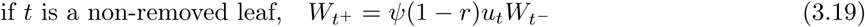

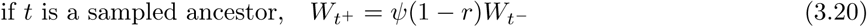

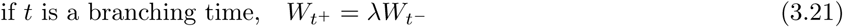

Combining these updates with equation (3.17) leads to the following proposition.

#### Proposition 3.1.

*At any time t across epoch* (*t*_*h*_, *t*_*h*+1_), *considering that we observed so far –i.e. on* (0, *t*_*h*+1_) *– v sampled ancestors, w removed leaves at times t*_*j*_ ∈ *𝒲, x branching events at times t*_*j*_ ∈ *𝒳, y non-removed leaves at times t*_*j*_ ∈ *𝒴, we get*,

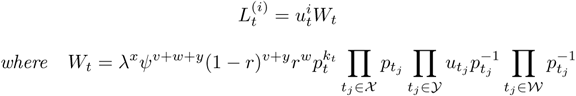

**Proof** We prove this proposition by induction across the epochs in Appendix D, using as the main arguments the equation updates (3.18) to (3.21), combined with equation (3.17).

Note that this proposition is very similar to what is presented in section 3 by Gupta et al. (2019). We nevertheless need to highlight two differences.

The first one is that we allow here for removal or not of the individual upon sampling, with a given probability *r*, whereas Gupta et al. (2019) considered that all individuals were removed upon sampling (*r* = 1), and Stadler (2010) considered that individuals were not removed upon sampling (*r* = 0).

The second difference concerns the underlying framework under which we derive our results. In Gupta et al. (2019), individuals where distinguishable (say, each one is assigned a number and they can be ordered), whereas in the present paper they are not. When individuals are ordered, the probability density 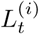 is changed by a factor 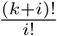, which is the number of ways we can arrange *k* + *i* elements in a list of size *k*, i.e. the number of ordered configurations of hidden individuals.

Note that, when reaching the root of the tree, it reduces to a very similar formula for the probability density of *𝒯* because *i* = 0 and *k* = 1. We summarize this as the following corollary.

#### Corollary 3.2.

*When ω* = 0, *the probability density of a reconstructed tree T with v sampled ancestors, w removed leaves at times t*_*j*_ ∈ *𝒲, y non-removed leaves at times t*_*j*_ ∈ *𝒴, and branching events at times t*_*j*_ ∈ *𝒳, is*

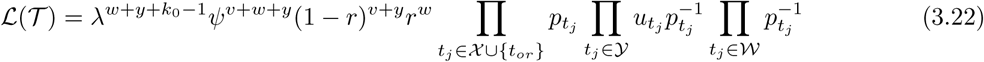

**Proof** It directly follows from Proposition 3.1, by noting that 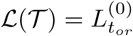. Note also that a rooted binary tree with *w* + *y* + *k*_0_ leaves shows necessarily *x* = *w* + *y* + *k*_0_ − 1 branching times.

### 3.3. Special case r = 1

When *r* = 1, only three kinds of punctual events, corresponding to updates (3.11), (3.14) and (3.16) need to be taken into account. Because the number of unsampled individuals *i* goes into formula (3.14), the simple expression 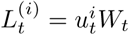 cannot be considered anymore, and one needs to find another expression. This has already been done in Gupta et al. (2019) and we only need to adapt here their result to our slightly different framework. For this we define the distinguishable version of the probability 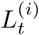 as

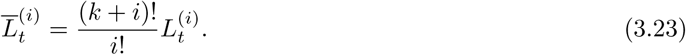

We now derive the Master equation for 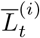. Multiplying both sides of (3.10) by 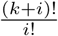 we obtain

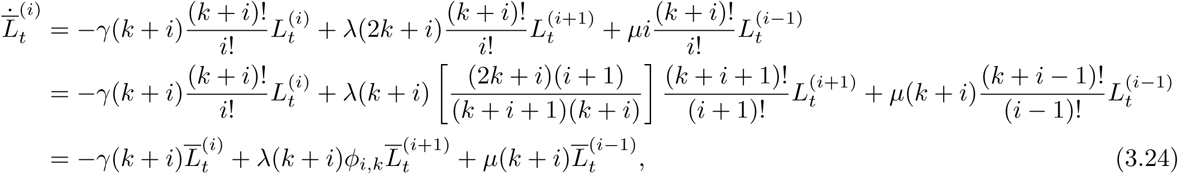

where

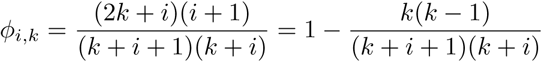

is the probability that a coalescing pair of randomly chosen lineages (from (*k* + *i* + 1) total lineages) does not consist of two sampled lineages. This shows that 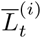 satisfies the Master equation (3.24) across any epoch. One can see that at punctual events the transition conditions (3.11) and (3.16) hold for 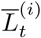 for *ψ*-sampling and branching events respectively. Moreover at *ω*-sampling events the transition condition (3.14) transforms to

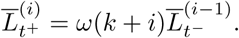

With these transition conditions and initial condition 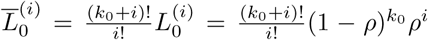, the Master equation (3.24) was solved explicitly in Gupta et al. (2019) and the solution is of the form

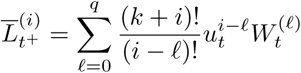

where *q* is the number of *ω*-sampling events in the time-interval [0, *t*) and the *q* + 1-dimensional time-varying vector 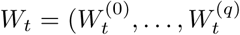 can be analytically computed following the approach in Gupta et al. (2019).

Therefore from (3.23) we state the following proposition.

#### Proposition 3.3.

*At any time t, we can compute the* 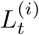 *values as*

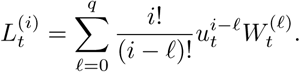

*where W*_*t*_ *is a q dimensional time-varying vector which can be computed following Algorithm 2 in Gupta et al. (2019)*.

Note that when there is no *ω*-sampling, then *q* = 0 for all times and 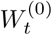 is the same as *W*_*t*_ defined in the previous section.

This ends our section on the computation of *L*_*t*_. It thus remains to (i) present a way to compute *M*_*t*_ and (ii)combine *L*_*t*_ and *M*_*t*_ to get the target distribution *K*_*t*_ at any time *t*. We do this in turn in the next two sections.

## 4. The joint density of observations above and past population size

Recall that we are now interested in computing,∀*i* ∈ ℕ, 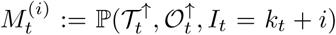. We start by presenting the Master equation satisfied by *M*_*t*_, before turning to its resolution for specific parameter sets. The approach is very similar to the one presented in the previous section to compute *L*_*t*_, with the slight difference that we will need to traverse the tree forward in time instead of backward in time.

### 4.1. Master equation driving M_t_

At the time of origin of the process *t*_*or*_, we only observe one starting lineage in 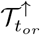. This provides us with the following initialization condition on *M*,

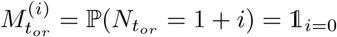

We then derive the Master equation driving the evolution of *M*_*t*_ across an epoch on which the number of observed lineages is fixed and equal to *k*. Suppose we know *M* _*t*_, and we observe no punctual event on the infinitesimal time interval (*t* − *δt, t*). Unobservable events have already been illustrated in Figure 2. It allows us to get

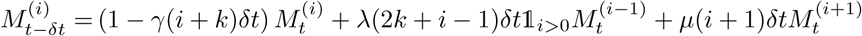

Letting *δt* → 0 and combining this with the initialization, we get the following set of ODEs driving the evolution of *M*_*t*_,

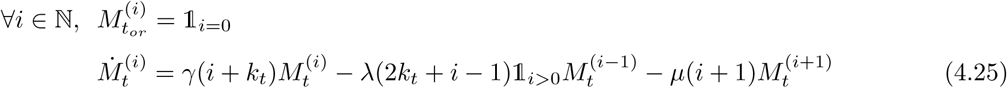

Last, we need to take into account the evolution of *M*_*t*_ at punctual events. Again, there are 6 types of punctual events that we can come across at time *t* in the past, listed below and illustrated in Figure 3. We denote *M*_*t*_− the probability just after (i.e. below) the punctual event and *M*_*t*_+ the probability immediately before (i.e. up). Because we are here deriving *M*_*t*_ forward in time, one needs to carefully note differences with results derived in section 3 relating to the number of lineages before and after the event. We can indeed find the same punctual events, namely,

1. a leaf of 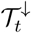, labeled as removed. This is a *ψ*-sampling with removal event for which the number of sampled lineages decreases by one and the number of unsampled lineages remains unchanged. This gives,

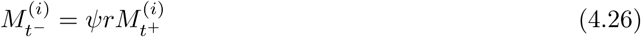
2. a leaf of 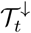, labeled as non-removed. This is a ψ-sampling without removal event for which one sampled lineages becomes unsampled. This gives,

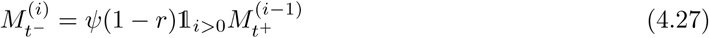
3. a sampled ancestor along a branch of 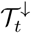, necessarily labeled as non-removed. This is a *ψ*-sampling without removal event which does not affect the number of lineages. It gives,

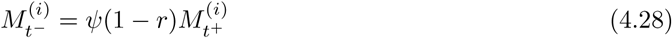
4. an occurrence in 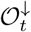, labeled as removed. This is a *ω*-sampling with removal event, for which the number of unsampled lineages decreases by one. This gives,

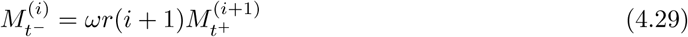
5. an occurrence in 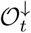, labeled as non-removed. This is a *ω*-sampling without removal event which does not affect the number of lineages. It gives,

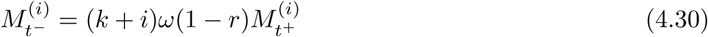
6. a branching event between two branches of 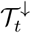.

This is a *λ*-event increasing the number of sampled lineages by one. This gives,

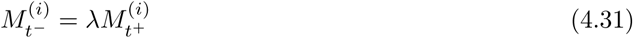

Finally, upon reaching present time 0, one needs to take into account the *ρ*-sampling, leading to the following update,

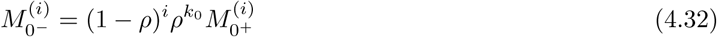

As already exposed for *L*_*t*_, we can build a similar algorithm to compute *M*_*t*_ in the general case, relying on a numerical ODE solver for approximating equation (4.25). As for algorithm 1, a slight variation of this algorithm would allow one to compute an estimate of the probability density of (*𝒯,𝒪*). Note that this strategy is very close to what has been used to compute the probability density of a reconstructed tree under a logistic birth-death process (Etienne et al., 2012).

In the remainder of this section, we derive analytical results to avoid resorting to a numerical ODE solver.

### 4.2. The corresponding probability generating function

We introduce now the probability generating function corresponding to the density *M*_*t*_, which will prove useful to get analytical results.

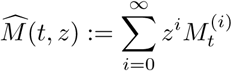

The initial condition on *M* translates in 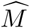 as,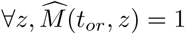. The ODE (4.25) furthermore translates as the following partial differential equation (PDE),

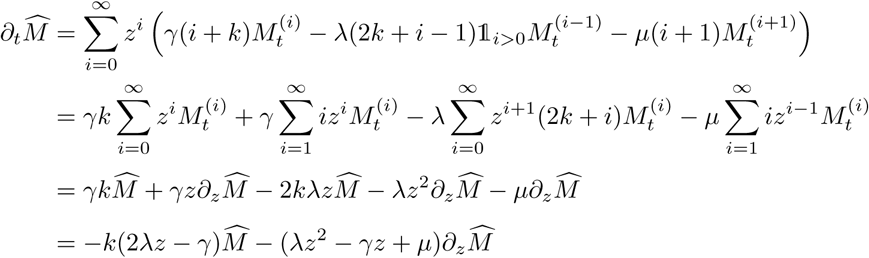

Our target probability generating function 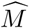 is thus the solution of the following PDE problem across a given epoch (*t*_*h*−1_, *t*_*h*_), on which the number of observed lineages remains constant and equal to *k*,

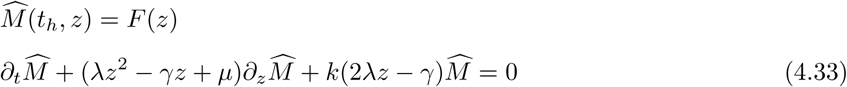

Solving this PDE problem allows us to come with an analytical expression of 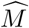 any time across an epoch, provided we now the expression of 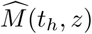 at the end of the epoch.

#### Proposition 4.1.

*The solution to the PDE problem (4.33) is given by*

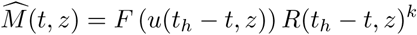

*where we introduce R*(*t, z*) = *p*(*t, z*)*/*(1 − *z*) *to ease notation*.

**Proof** We used the method of characteristics, see derivations in Appendix B.

Between epochs, one must also update 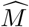 according to punctual events taking place. Previously presented updates of *M* (equations (4.26) to (4.31)) translate as the following for 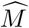,

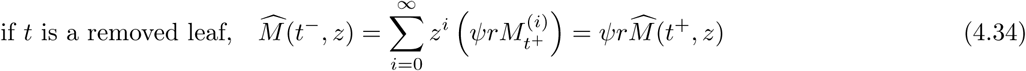

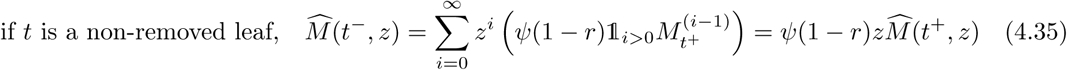

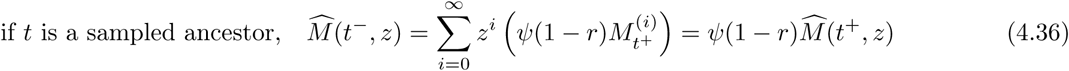

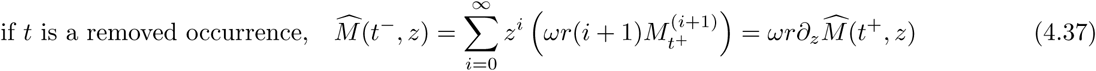

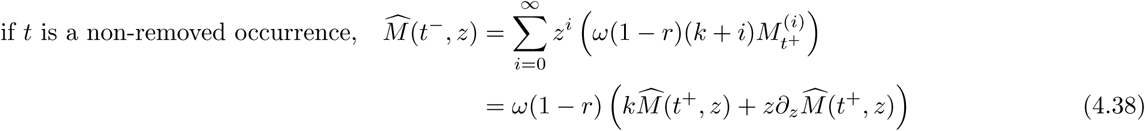

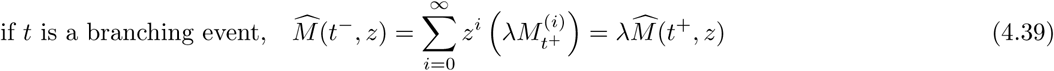

If we are interested in the distribution at some point, we can thus start the formula at *t*_*or*_ with *F* (*z*) = 1, and then iteratively alternate between the updates at punctual events and the use of Proposition 4.1 over each epoch. When reaching present time 0, the step of *ρ*-sampling expressed in equation (4.32) moreover translates here as,

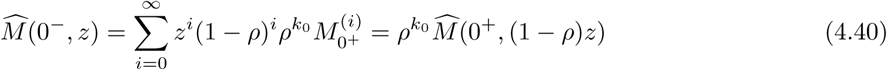

While this procedure in theory allows us to get the analytical formula of 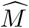 at any time, updates (4.37) and (4.38) require differentiating the probability generating function, greatly complicating the expression of the function after a few occurrences. When *ω* = 0, these two updates disappear and a nice recursion leads to a closed-form formula that we will detail in Proposition 4.3.

We also implemented this procedure in a programming language able to deal with symbolic calculus, but were not able to provide an implementation efficient enough to apply to standard datasets in the field. Instead, when *ω*≠ 0, we suggest another strategy for computing the 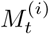’s, by approximating 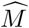 across punctual events by a polynomial of order *N*, while still relying on Proposition 4.1 to drive the evolution of the probability generating function between events. This is a more efficient alternative to numerically solving the ODE system. We only need to derive the expression of the generating function at punctual events as given in the following proposition 4.2.

#### Proposition 4.2.

*The derivatives in z* = 0 *of a generative function which expression is*

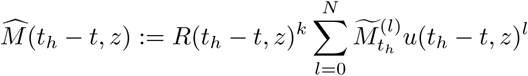

*can be numerically computed using the formula*

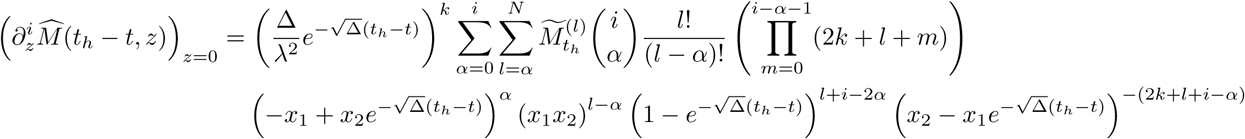

**Proof** The derivation is detailed in Appendix C.1.

This derivation is at the heart of Algorithm 2, for it allows to drive the evolution of the 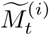’s through each epoch, as well as at times when we want to record them.

We will refer to Algorithm 2’ as the slight variation of this algorithm aimed at computing the density of (*𝒯*, *𝒪*). No set of time points (*τ*_*j*_) is required, and the values of 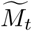 are not recorded through time (i.e. matrix *B′* disappears). Instead, when reaching *t*_*h*_ = *t*_0_ we simply return 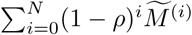

#### Algorithm 2 Computes a numerical approximation of *M*_*t*_ for a specific set of times

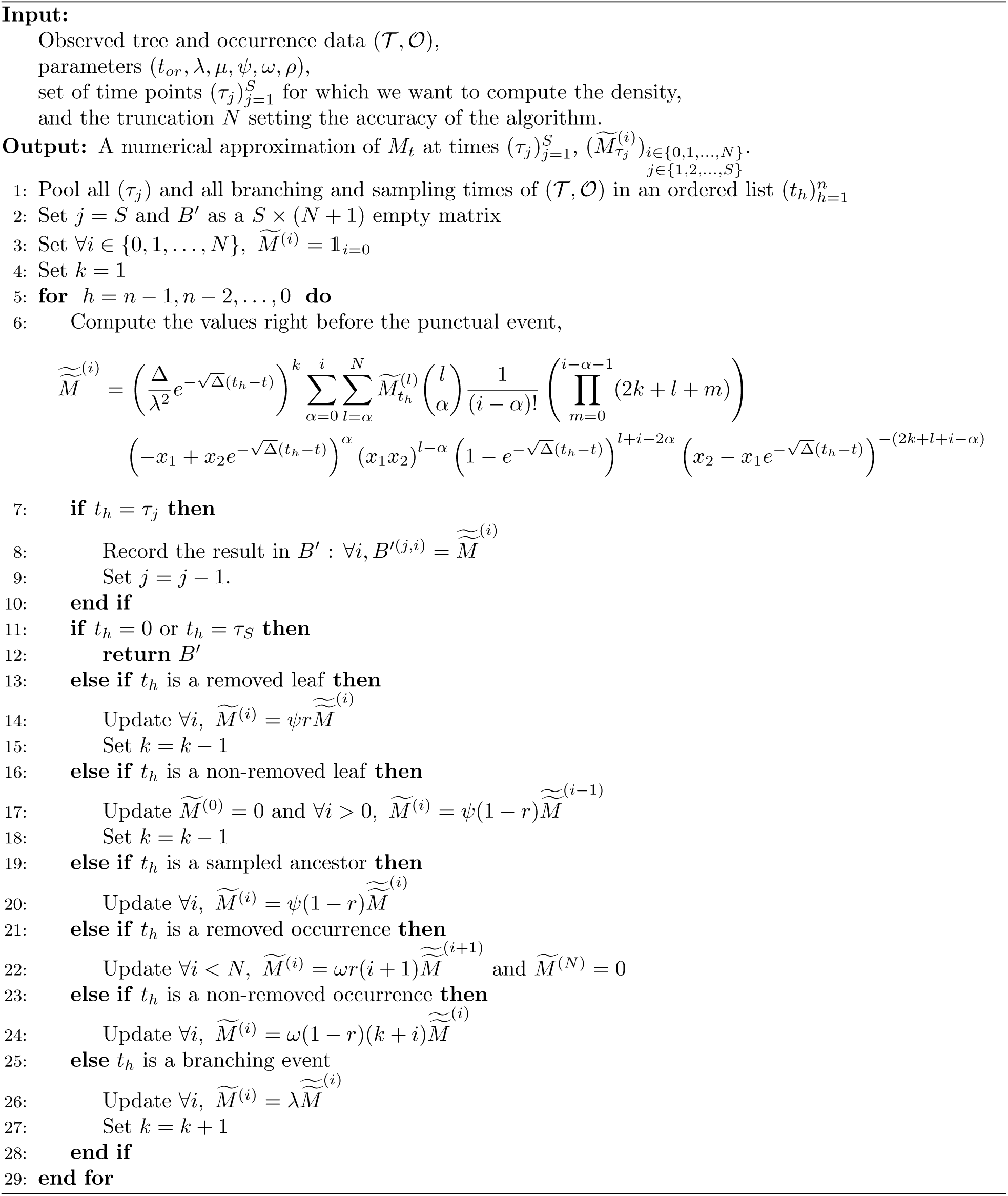

One may wonder why we introduced a probability generating function in this section and not in the previous one to fasten Algorithm 1. It turns out that we tried to follow the same approach to derive *L*_*t*_, as described in Appendix E, but it leads to another PDE problem that will require further work to be solved.

### 4.3. Special case ω = 0

This corresponds to the special case leading to the observation of *𝒪* = ø. In that case, a nice recursion leads to a closed-form formula for 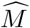.

#### Proposition 4.3.

*At any time t, considering that we observed so far –i.e. on* (*t, t*_*or*_) *– v sampled ancestors, w removed leaves at times t*_*j*_ ∈ *𝒲, x branching events at times t*_*j*_ ∈ *𝒳, y non-removed leaves at times t*_*j*_ ∈ *𝒴, we get*,

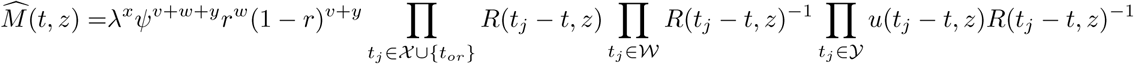

**Proof** We prove this result by induction across the epochs of *𝒯* in Appendix D, using as the main arguments the update equations (4.34), (4.35), (4.36), (4.39), combined with Proposition 4.1 driving the evolution across an epoch.

As a simple corollary of this result, when *t*_*h*_ = 0 is the present, we get back the same probability density formula of *𝒯* as provided, e.g. in theorem 3.5 in Stadler (2010) (when *r* = 0), in section 3 in Gupta et al. (2019) (when *r* = 1), or in our previous corollary 3.2.

**Proof** Proposition 4.3 offers yet another proof of corollary 3.2 by noting that

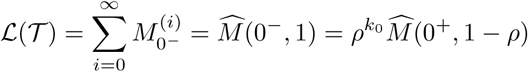

where the last equality follows from equation (4.40) taking into account the *ρ*-sampling at present.

When *ω* = 0, Proposition 4.3 also offers an alternative to Algorithm 2 for deriving *M*_*t*_. Indeed, resorting to the probability generating function to get back the probability density, one can get the following corollary.

#### Corollary 4.4.

*At any time t, considering that we observed so far –i.e. on* (*t, t*_*or*_) *– v sampled ancestors, w removed leaves at times t*_*j*_ ∈ *𝒲, x branching events at times t*_*j*_ ∈ *𝒳, y non-removed leaves at times t*_*j*_ ∈ *𝒴, we can compute* 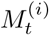 *using the following recursion*,

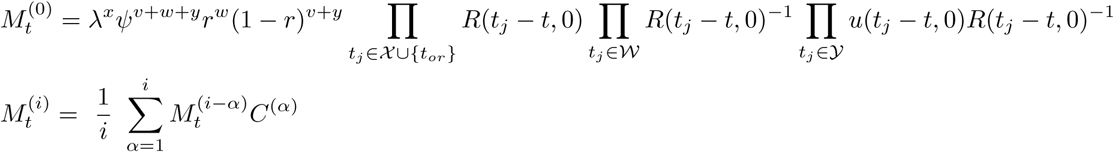

*where we define*

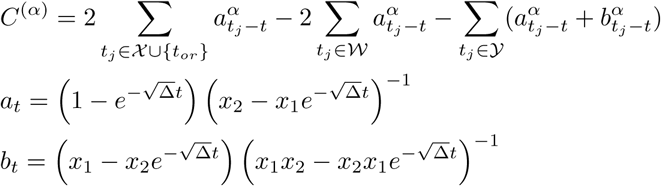

**Proof** The probability density 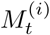 can be found back by taking

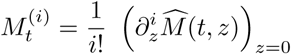

The result follows from the derivation of these derivatives in Appendix C.2.

This special case ends the section. In the next one, we will combine results from sections 3 and 4 and use our ability to compute *L*_*t*_ and *M*_*t*_ to compute *K*_*t*_, the probability distribution of the population size given (*𝒯*, *𝒪*).

## 5. The distribution of past population size conditioned on observations

### 5.1. The distribution at fixed times

In section 3, we exposed how to compute *L*_*t*_, the probability density of the observations below time *t* conditioned on the population size. This relies either on Algorithm 1 in the general case, or on the more optimized Proposition 3.1 in case *ω* = 0, or Proposition 3.3 in case *r* = 1.

In section 4, we exposed how to compute *M*_*t*_, the probability density of the observations above time *t* and the population size. This relies either on Algorithm 2 in the general case, or on the more optimized Corollary 4.4 when *ω* = 0.

We now combine our ability to compute *L*_*t*_ and *M*_*t*_ to derive the probability distribution of the population size given (*𝒯,𝒪*). Recall from the first section that,

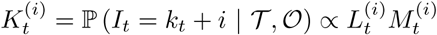

Provided we have stored numerical values 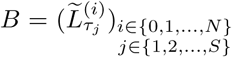 and 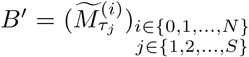 for a set of time points 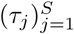, it only remains to compute

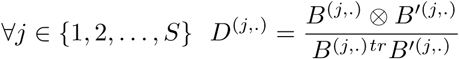

where *⊗* refers to the Hadamard product, and ^*tr*^ is the transpose of the vector.

Depending on the parameter space that one wants to consider, it thus remains to arrange pieces stemming from the previous sections. We provide a flowchart in Figure 4 to guide the reader to chose the most efficient path.

**Figure 4:**
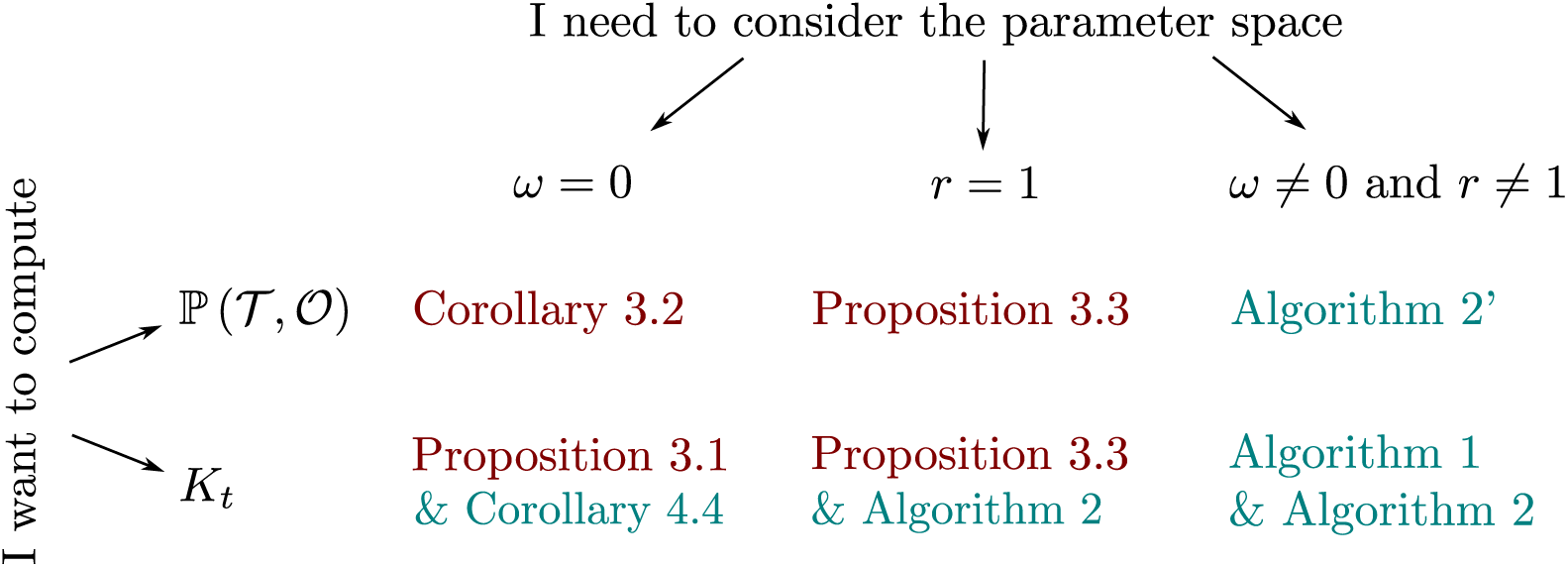
The most efficient results depending on the parameter space considered. In red, results already described in Stadler (2010) and Gupta et al. (2019). In blue, the new contribution of this manuscript.

### 5.2. Generator of trajectories

The previous result gives us the distribution of the population size at any time in the past, but does not state anything about population size trajectories. We provide now an approximate way of simulating population size trajectories conditioned on (*𝒯*, *𝒪*).

Indeed, recall we have,

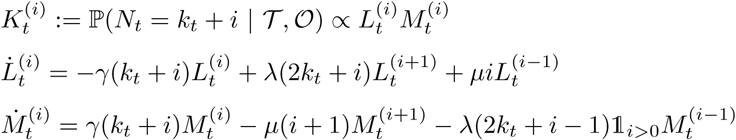

We thus get,

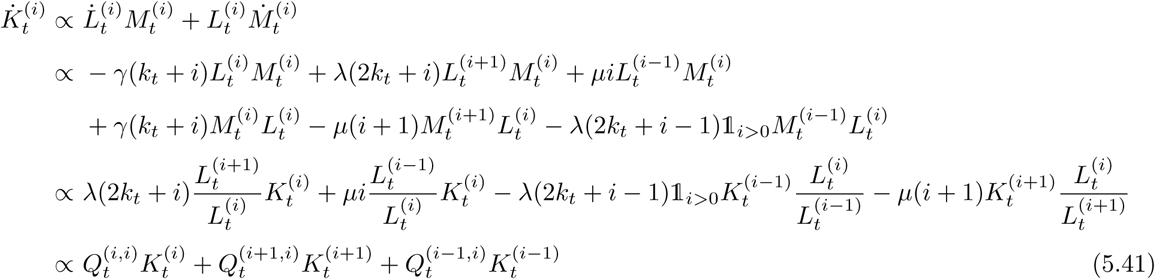

Where we introduced in the last line the following notation,

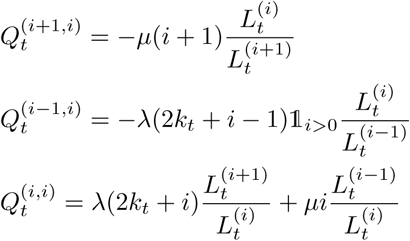

Using these, we see that 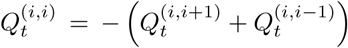. This allows us to draw trajectories of the number of ancestors in the past as a continuous in time Markov process with the (inhomogeneous) rates *Q*_*t*_ written above.

Remark that we could equally write these ODE coefficients using the 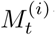’s, depending on which traversal we prefer to perform prior to the simulation. This gives,

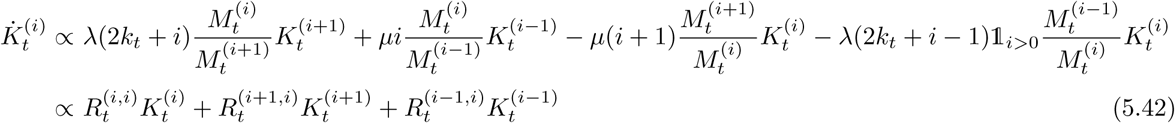

where we introduced in the last line the following notation,

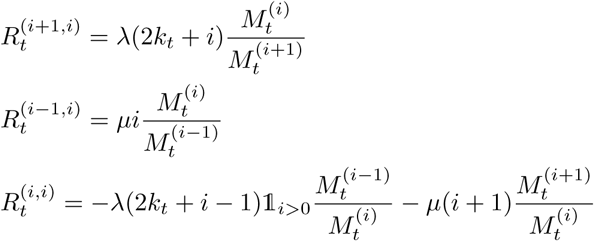

This is a standard result for Markov chains that are conditioned on a final state, and the shape of the newly derived transition kernel is called a Doob’s transform. Note that these transitions symplify for special cases when we have an analytical expression of either 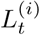 or 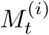

### 5.3. Numerical implementation

Results of this paper have been implemented numerically and the code is freely available on GitLab: *https://gitlab.com/MMarc/popsize-distribution/*.

We used the numerical implementation to verify the correctness of the results in several ways:

1. We verified that the values of the probability density of (*𝒯*, *𝒪*) computed using *L*_*t*_ and *M*_*t*_ (algorithms 1’ and 2’) were equivalent to values computed using already known formulas when *ω* = 0 (Stadler,2010) or when *r* = 1 (Gupta et al., 2019). See result in Figure 5AB.
2. We verified that the values of the probability density of (*𝒯*, *𝒪*) computed using *L*_*t*_ or *M*_*t*_ (algorithms 1’ and 2’) were identical on examples for which no previous formula was known. See result in Figure 5C.
3. We assessed the distribution of the population size against the only numerical method performing the same goal, the particle filtering developed in Vaughan et al. (2019). We compared values of a few quantiles computed using the two methods, see result in Figure 5DEF).

We also illustrate in Figure 6 our target distribution *K*_*t*_ of the past population size conditioned on (*𝒯*, *𝒪*), on a few simulated examples.

**Figure 5:**
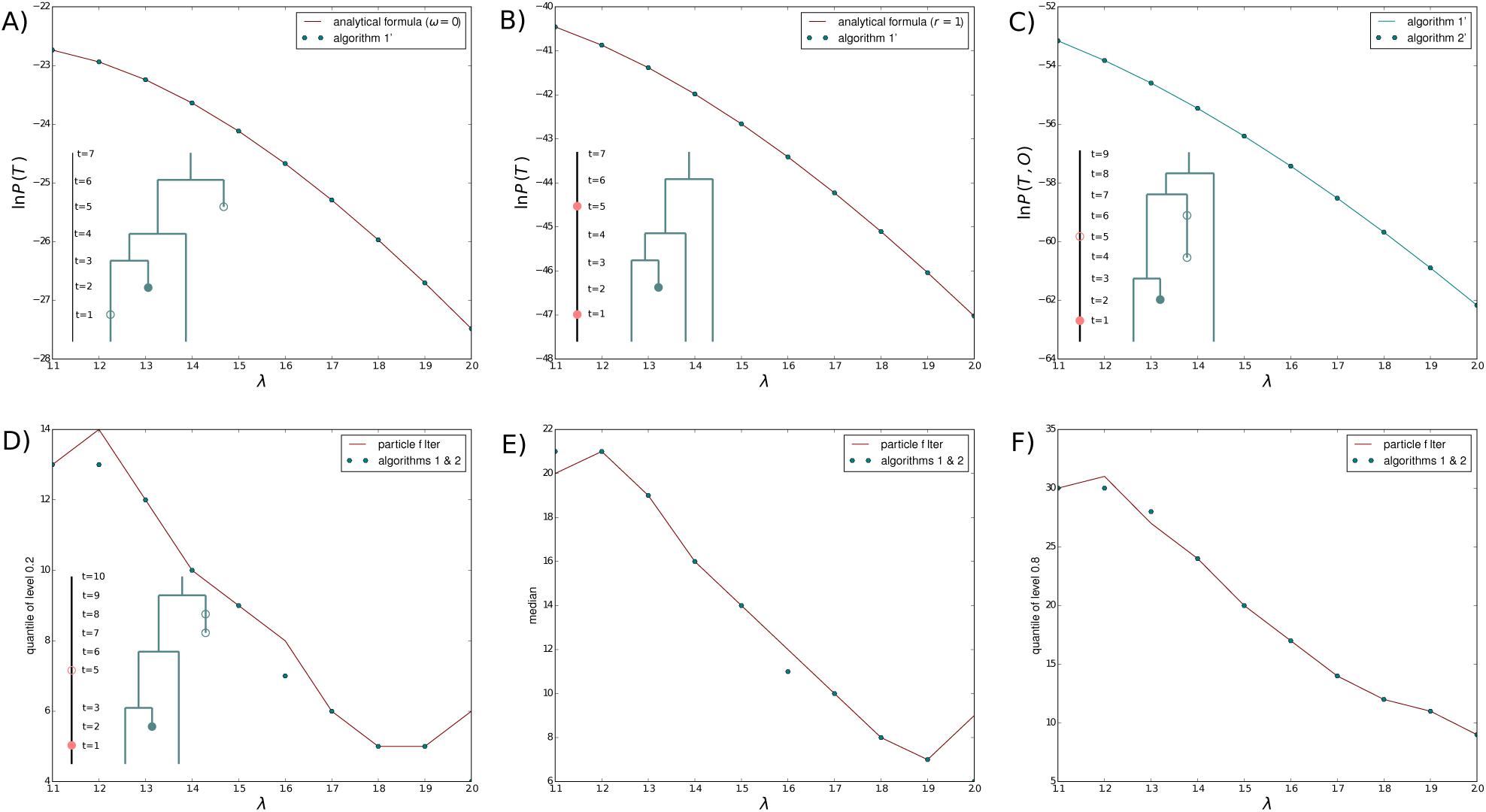
Assessment of the accuracy of the methods presented in this paper, on toy datasets. First row, probability density of data, A) against known analytical formula when *ω* = 0 and (*µ, ρ, ψ, r*) = (1, 0.5, 0.3, 0.2); B) against known analytical formula when *r* = 1 and (*µ, ρ, ψ, ω*) = (1, 0.5, 0.3, 0.6); C) obtained using algorithms 1’ or 2’ otherwise, with (*µ, ρ, ψ, r, ω*) = (1, 0.5, 0.3, 0.2, 0.6). Second row, quantiles of the population size distribution, against the particle filter in Vaughan et al. (2019), with parameters (*µ, ρ, ψ, r, ω*) = (1, 0.1, 0.001, 0.5, 0.001). D) quantile of level 0.2; E) median; F) quantile of level 0.8.

**Figure 6:**
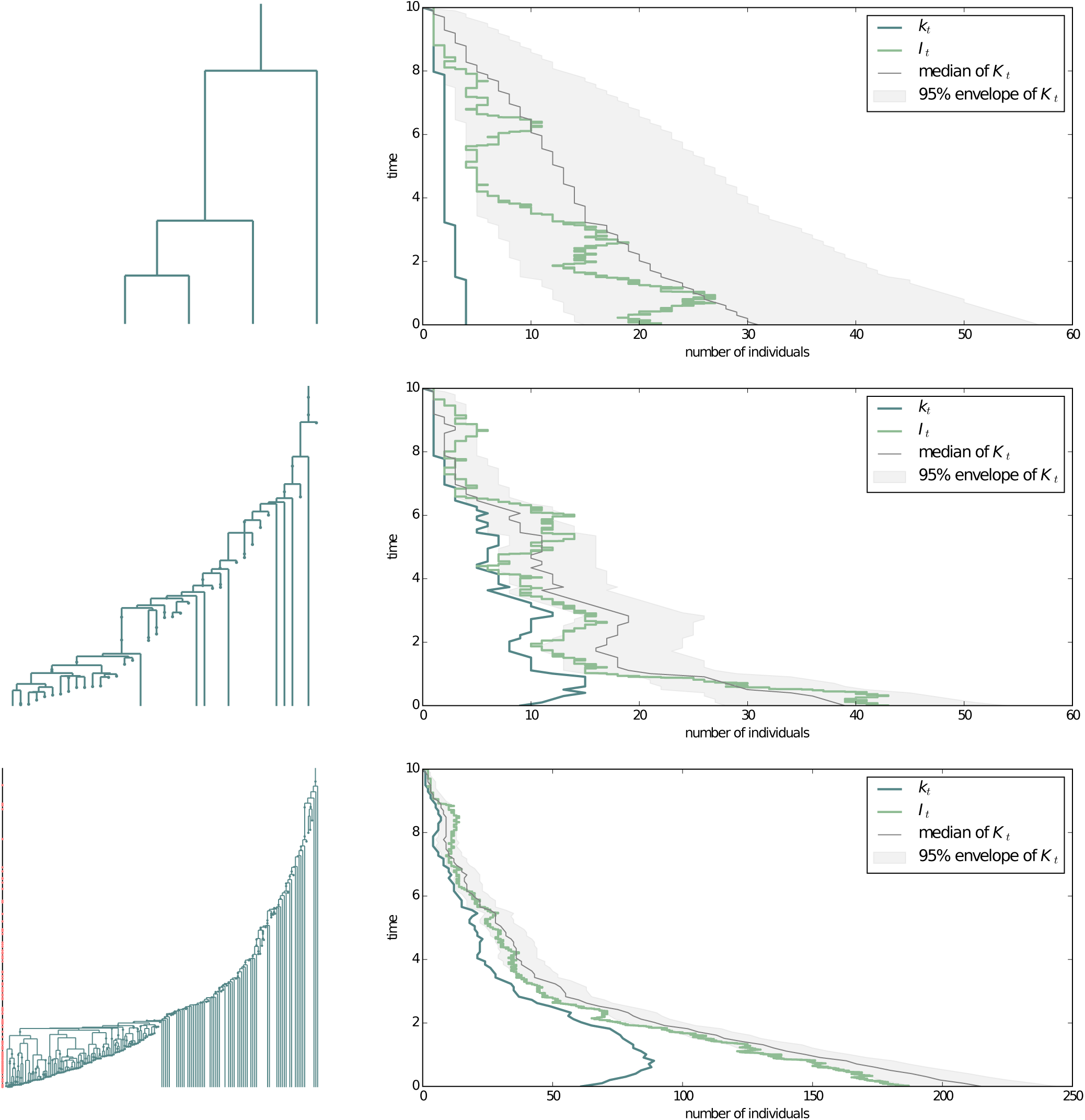
Hints of the true population size (in green) under three different processes. A) A homogeneous birth-death with *ρ*-sampling at present; B) A homogeneous birth-death with *ρ*-sampling at present and *ψ*-sampling through time; C) A homogeneous birth-death process with *ρ*-, *ψ*- and *ω*-sampling.

## 6. Discussion

The results we have derived in this paper fit into two main categories.

The first category concerns results allowing one to compute the probability density *ℒ* of (*𝒯*, *𝒯*). While results allowing to compute *ℒ* when *ω* = 0 or *r* = 1 were already known, our two algorithms 1’ and 2’ have the potential to improve the computation time of *L* when *ω* ≠ 0 and *r* ≠ 1.

The second category concerns our main results, allowing one to compute the probability distribution of the population size in the past and to generate population size trajectories conditioned on (*𝒯*, *𝒪*) (section 5). These constitute a dramatically more efficient alternative to the Monte-Carlo particle filtering algorithm developed by Vaughan et al. (2019) to perform the same task.

We discuss first how these results can be used in the context of current phylodynamics, before turning to future extensions.

### 6.1. Use for inference purposes

When applying phylodynamics methods to empirical data, it is common practice in the field to condition the model generating the phylogeny on either (i) the observation of at least one individual at present, descending from time *t*_*or*_, an event that we call 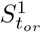; or (ii) the observation of at least two individuals at present, descending from time *t*_*mrca*_, an event that we call 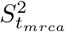, where *mrca* stands for *most recent common ancestor*. Indeed, we can study a dataset precisely because it survived to the present.

The probability density of (*𝒯*, *𝒪*) needs be conditioned on 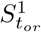 or 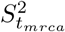 when used in empirical applications, which can be done by dividing *ℒ* by, respectively, 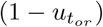 and 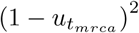 However, since 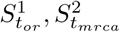 are always satisfied on empirical data, and we already condition on the observation of empirical data, nothing more needs to be done to use the probability distribution *K*_*t*_ to retrieve past population sizes.

If the tree is directly fixed and observed, results from the first category can be used to perform parameter inferences in a maximum likelihood framework. Taken as a function of the parameters, the probability density of (*𝒯*, *𝒪*) is called the likelihood, and can be numerically optimized to look for parameters maximizing its value (i.e. maximum likelihood estimators).

If one also knows the true parameter values or has inferred estimates of those, results from the second category can also be used to compute the distribution of the population size in the past. Any quantile of interest can then be numerically computed from this distribution. As an example, we chose to represent on Figure 6 the 95% envelope of the distribution, comprised between the level 2.5% and the level 97.5% quantiles. Alternatively, if one needs a point estimate of the population size, the median of the distribution is a good candidate.

If the tree is not directly observed, but sequencing data or any other set of phenotypic measurements, called *A*, is available for the set of *ψ*-sampled individuals, then one might consider analyzing those in a Bayesian framework. For instance, consider a model describing the evolution of molecular sequences or phenotypic measurements along a fixed t ree i s c hosen, r elying o n p arameters t hat w e c all *θ*. We a re then interested in quantifying the posterior distribution

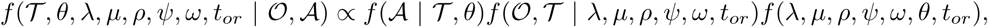

with *f* (*λ, µ, ρ, ψ, ω, θ, t*_*or*_) being the prior distribution on the model parameters. Standard MCMC algorithms can be used to sample from this posterior distribution. Results from the second category can then be used and applied to a sample from the posterior distribution to integrate the distribution of the population size over the posterior distribution of trees and parameters values.

In addition, the distribution *K*_*t*_ of population sizes (or a trajectory sampled from this distribution using the results of section 5.2) can be computed directly for each sampled combination of tree and parameters, and thus used to compute the full joint posterior of both population size trajectories, trees and model parameters. While this is similar to the capability of the method of Vaughan et al. (2019), we expect the method presented here to be far more computationally efficient as it cuts out the necessity of simulating particle trajectories within the MCMC step. (The trajectory specific c alculations o r s imulations i n our method need only be done *after* the MCMC algorithm is complete, and only for a subsample of the full MCMC chain states.)

### 6.2. Future extensions

This process lends itself well for various biologically realistic extensions to allow closer fit t o empirical data in a variety of situations.

The first extension that we envision is to relax the assumption of rate homogeneity and instead work with time-varying rates. This has already been considered in different studies relying on birth-death processes, either with exponentially varying functions (Morlon et al., 2011) or with piecewise constant rates (a model dubbed as *skyline birth-death process*, see Stadler et al., 2013; Gavryushkina et al., 2016). Because most of what we wrote can be straightforwardly adapted to such a framework, this would not require much theoretical work. However, the challenge would be to do so without overfitting the data.

Another popular extension that has been described in the literature on birth-death processes for phylo-dynamics is to consider multi-type birth-death processes (Maddison et al., 2007). Each individual is assigned a type, which impacts its propensity to give birth to other types. All sampling-related parameters can also be considered type-dependent. The challenge here boils down to dealing with an increase of dimensionality, where we would be interested in the joint distribution of subpopulation sizes. Recent work by Freyman and Höhna (2018) could first allow us to find the distribution of the types along sampled lineages in the tree. If we have *j* types, we would then be interested in

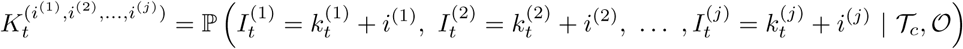

where all quantities are type-dependent analogs of quantities defined in the current paper, and *𝒯*_*c*_ would be a type-coloured tree, i.e. a tree where each sampled lineage is assigned a type. This extension is particularly interesting for epidemiological applications, when different populations of infected individuals, clustered according to some characteristic (e.g. patient behaviour or geography) might have very different dynamics (Stadler and Bonhoeffer, 2013).

Finally, we are very hopeful that this piece of work could be applied as well to density-dependent birth-death processes, also known as *logistic birth-death models*. Indeed, very similar ideas to the breadth-first forward and backward traversals as applied in algorithms 1’ and 2’ appear in the context of logistic birth-death models (Etienne et al., 2012; Leventhal et al., 2013). Preliminary results obtained by adapting our numerical algorithms to this framework are very encouraging, and we are currently in the process of deriving as much analytical results as we can to speed up the method. We are hoping to present this in a subsequent paper.

### 6.3. Conclusion

This manuscript presents a way to efficiently compute the distribution of the number of ancestors in a linear birth-death process, conditioned on the observation of a reconstructed tree and a record of occurrences through time. The method paves the way to the consideration of more complex demographic scenarios, with either time-dependent or density-dependent parameters. We hope that it will foster important research advances for unravelling demographic histories in epidemiology, macroevolution, and any other fields where birth-death processes form a relevant model framework.

## Acknowledgements

The authors are very grateful to Rachel Warnock for helpful comments on potential applications of the model.

## Appendix A. Solving well-known ODEs

### Appendix A.1. The extinction probability

We first deal with equation (2.1) governing *u*_*t*_, and start by studying the polynomial *λx*^2^ − *γx* + *µ*.

This polynomial has discriminant Δ = *γ*^2^ − 4*λµ >* (*λ* + *µ*)^2^ − 4*λµ ≥* (*λ* − *µ*)^2^ *>* 0. Roots are

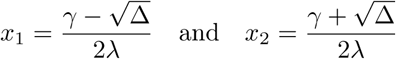

Moreover, we know that both roots are positive because 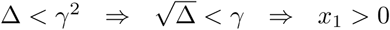.

On an interval including zero and where the polynomial remains positive (as (−∞, *x*_1_) for example), we can write,

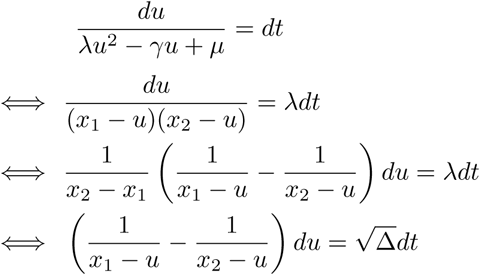

Integrating both sides between time 0 and *t*, we get

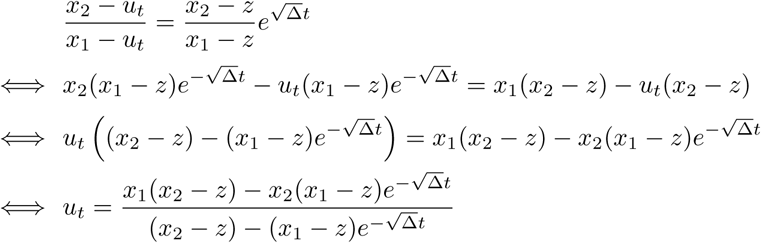

This is the result stated in equation (2.2). Note that this quantity is called *p*_0_(*t*) in Stadler (2010), or *E*(*t*) in Maddison et al. (2007).

### Appendix A.2. Probability to leave only one sampled descendent

We aim here to integrate a slight variation of equation (2.3) governing *p*_*t*_ when *k* = 1. The equation we are interested in is,

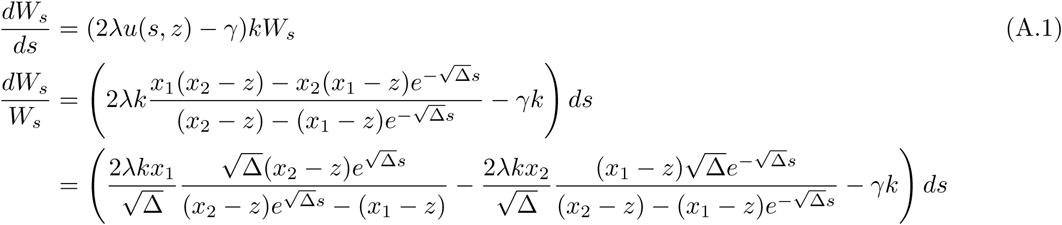

All these three terms can be integrated visually between some time *t*_*h*_ and *t*, leading to,

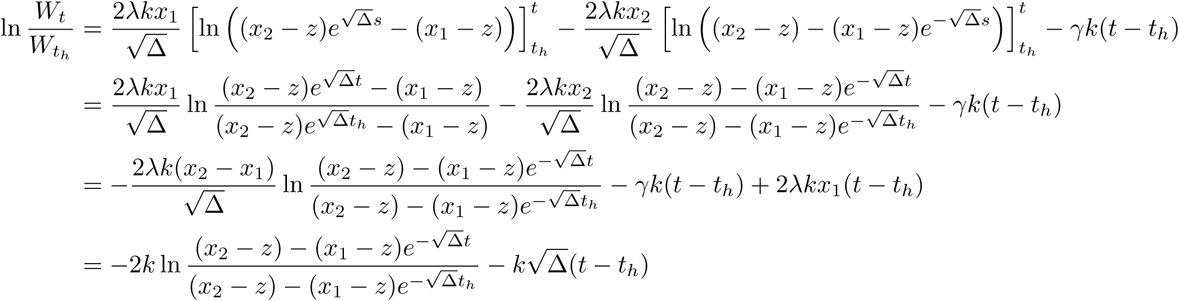

Leading to the final expression below

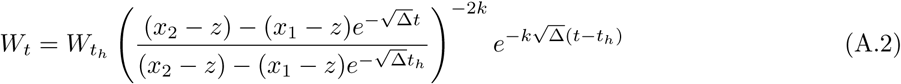

Note that the case *k* = 1, *t*_*h*_ = 0 and *W*_0_ = 1 − *z* corresponds to the probability *p*_*t*_ given as equation (2.4),

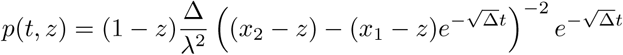

while the general case can be expressed using function *p* as

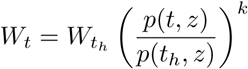

### Appendix A.3. A few useful properties

Solutions *u*(*t, z*) and *p*(*t, z*) to ODEs (2.1) and (2.3) satisfy two properties relying on the semi-group property of solutions of ODEs, namely,

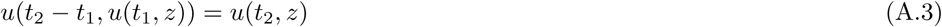

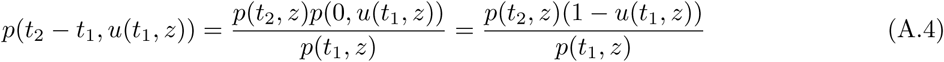

These two properties are useful in many calculations throughout this document, e.g.

- Solving the main PDE in Appendix B requires inverting *u*, using the first property with,

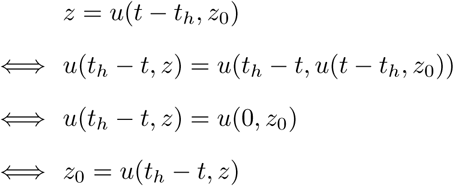
- The same Appendix section requires also composing function *p* and *u*, using

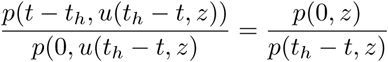
- In the proof of Proposition 4.3, we switch to the notation *R*(*t, z*) = *p*(*t, z*)*/*(1 − *z*) and again compose *R* and *u* in the same way,

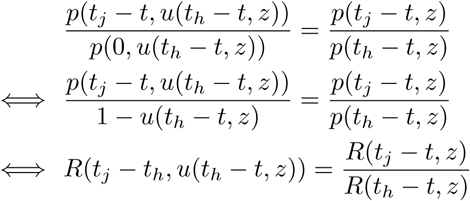

## Appendix B. Solving the main PDE

We aim now at finding an analytical solution for the following PDE, driving the evolution of 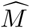 across a given epoch (*t*_*h*−1_, *t*_*h*_), on which the number of observed lineages remains constant and equal to *k*.

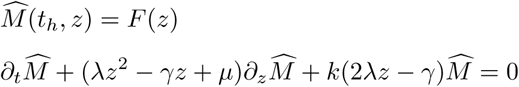

### Appendix B.1. Principle of the method of characteristics

This problem can be solved by the method of characteristics. We suppose that we can write 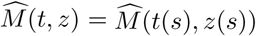 where functions *t* and *z* satisfy the ODEs,

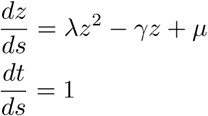

This way, the function 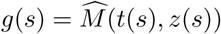 satisfies another ODE, that we will have to solve,

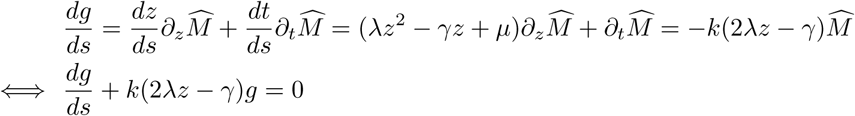

### Appendix B.2. Step 1, solve for t(s), z(s) and g(s)

We start by integrating *t*(*s*). We moreover fix that *t*(0) = *t*_*h*_, thus leading to *t*(*s*) = *t*_*h*_ + *s*.

We now turn to *z*, and remark that it satisfies previously studied ODE (2.1). Integrating between 0 and *s* leads to,

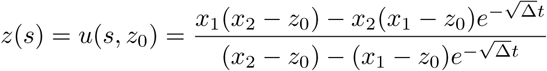

Last, *g* satisfies an ODE very similar to (A.1). Taking care of the minus sign, it leads to the following result,

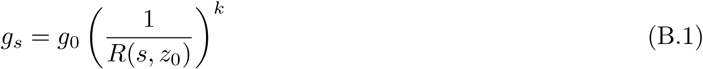

### Appendix B.3. Step 2, express 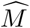 *back as a function of t, z*

We want to express our two unknown quantities *s* and *z*_0_ as functions of *t* and *z*.

On a first h and, w e g et e asily *s* = *t* − *t* _*h*_. W e m oreover c an s olve f or *z* _0_ i n t he f ollowing equation, remembering the semi-group property of *u*,

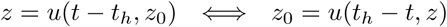

Substituting these into the previous expression (B.1) of *g*_*s*_ then leads to,

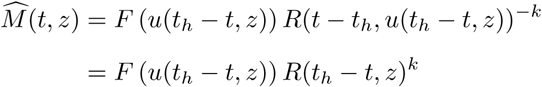

where the first to second equality relies on a property exposed in Appendix A.3. This gives us the final formula which is stated in Proposition 4.1.

## Appendix C. Some useful algebra

This section of the Appendix pools together all bits of algebra that are not really digestible, but are used in the main text.

### Appendix C.1. Deivative of 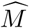

We first modify a bit the expression of the generating function,

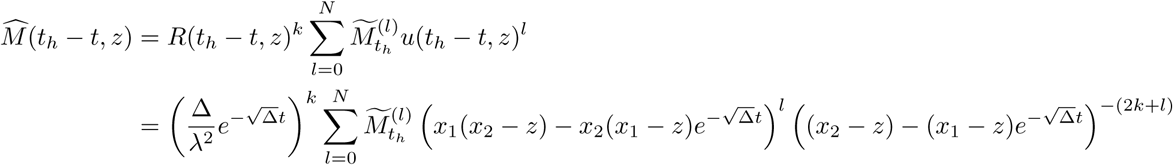

Applying Leibniz’s derivation rule to the product, we get,

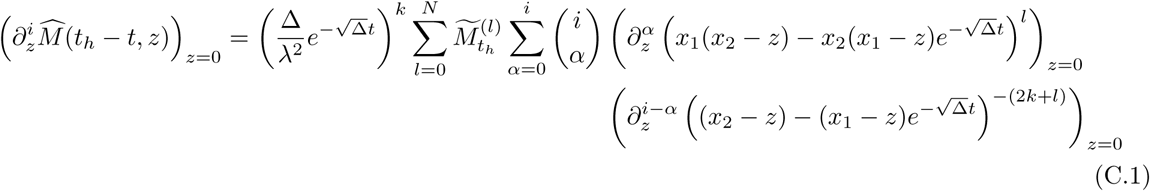

The first of the two derivatives in the sum can be computed as,

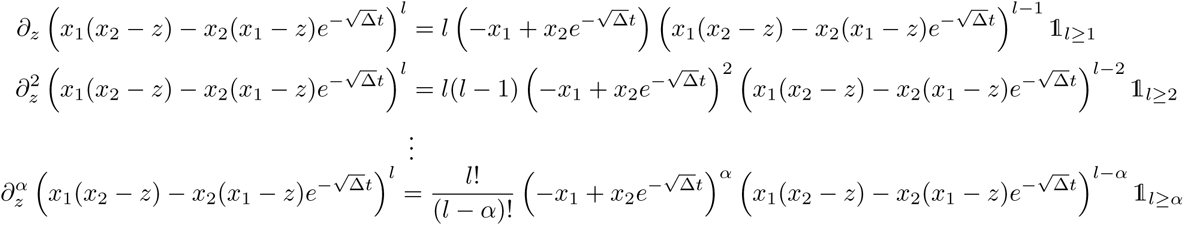

While the second gives us,

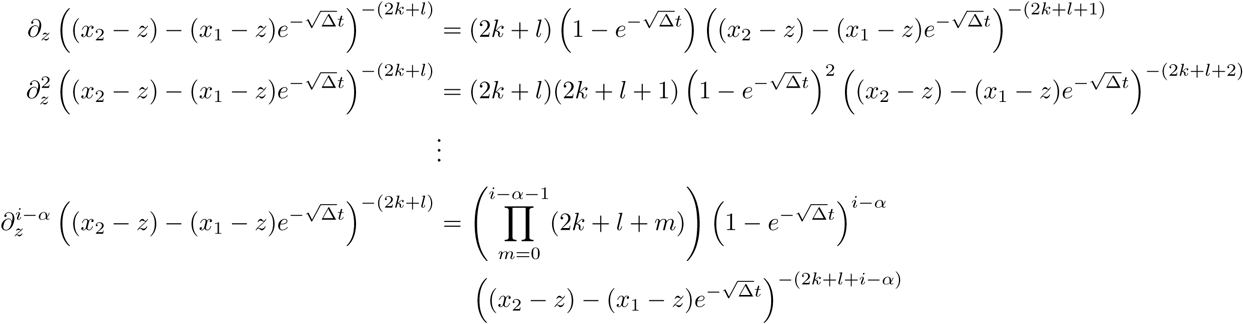

Applying these derivatives in *z* = 0 in equation (C.1) yields,

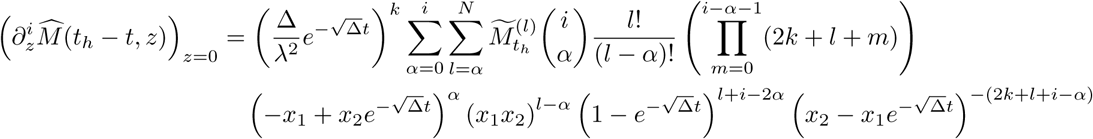

Which is the expression provided in Proposition (4.2).

### Appendix C.2. Derivatives of 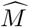 when ω = 0

We wish here to derive the 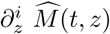 where function 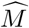 is as given in Proposition 4.3, i.e.

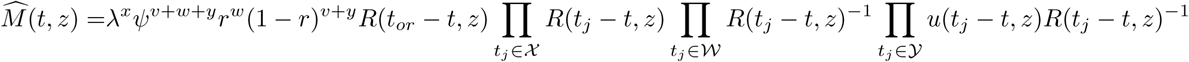

We take for simplicity the derivative of the logarithm of 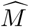 and express the derivatives of 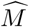 using these and Leibniz’s formula,

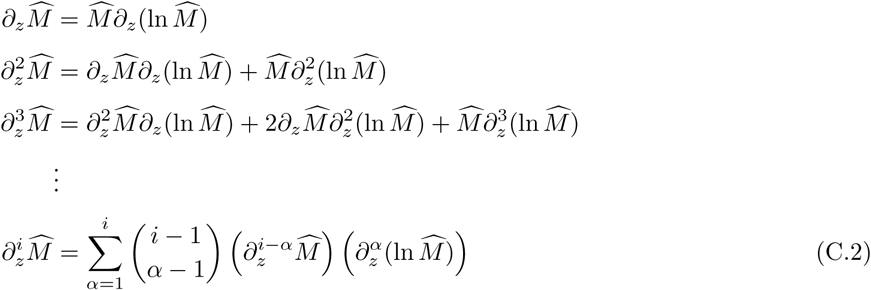

In order to compute the derivatives of ln 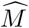, one needs to get the derivatives of ln *R*(*t, z*) and ln *u*(*t, z*). We have

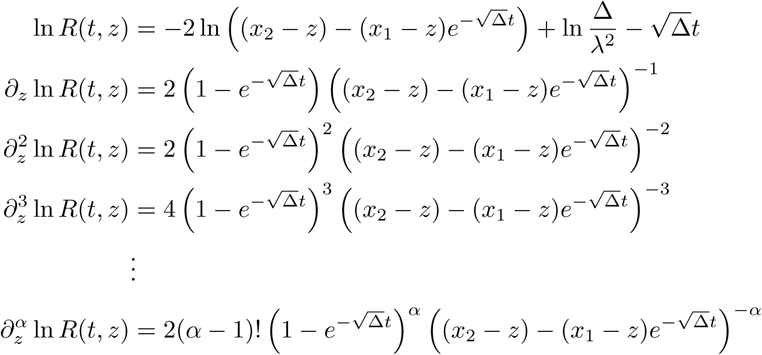

Finally taking the function in *z* = 0 leads to

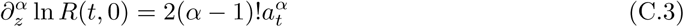

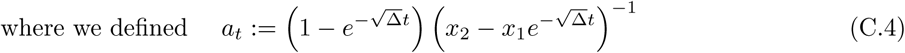

In the same way we get,

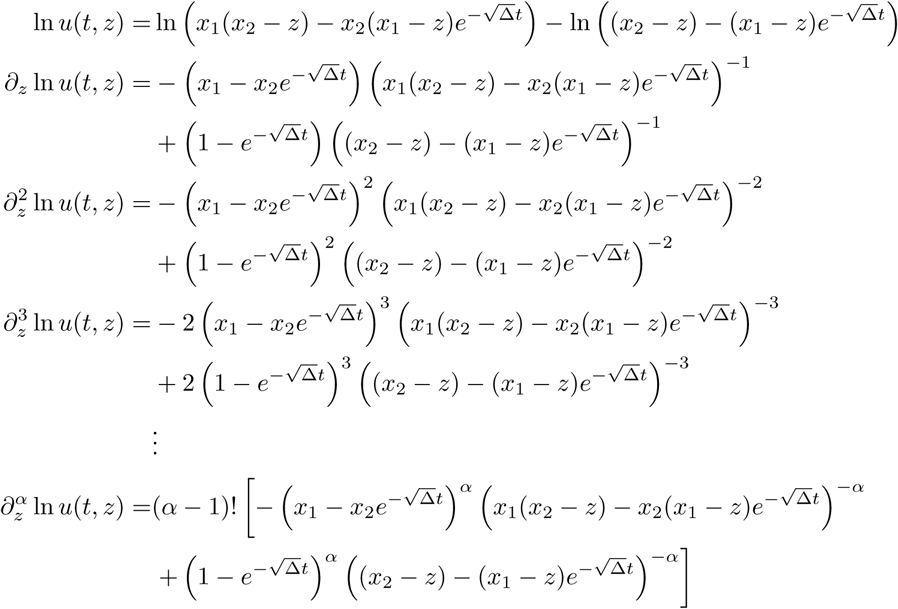

Here also, we are interested in the function in *z* = 0,

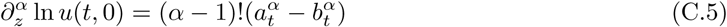

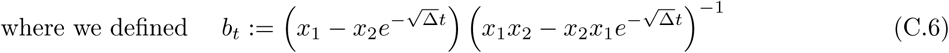

Last ingredient needed to write the derivative of ln 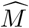, we get,

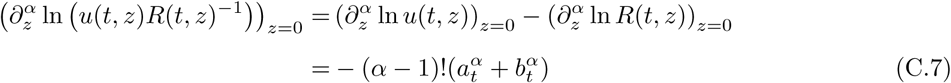

Finally, using equations C.4 and C.7, one can compute

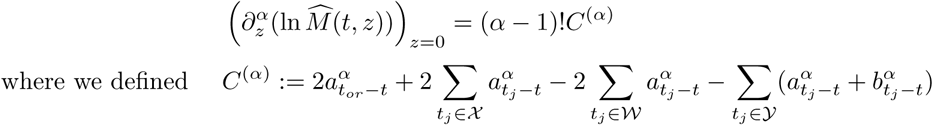

Plugging this into equation C.2 and noting that 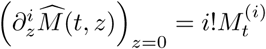, we get

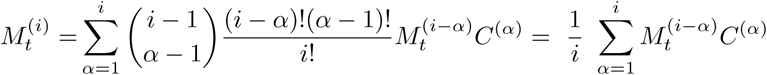

which is the result stated in corollary 4.4.

## Appendix D. Inductions across the epochs

### Appendix D.1. Proof of Proposition 3.1

We prove the proposition by induction across the epochs.

If we observe only the first epoch and the *k*_0_ leaves at present, then we get at any time *t* across the first epoch 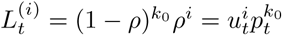, which satisfies Proposition 3.1.

Suppose we observed so far – i.e. on (0, *t*_*h*+1_) – *v* sampled ancestors, *w* removed leaves at times *t*_*j*_ ∈ *𝒲, x* branching events at times *t*_*j*_ ∈ *𝒯*, *𝒳* non-removed leaves at times *t*_*j*_ ∈ *𝒴*. And suppose that Proposition 3.1 is verified across epoch (*t*_*h*_, *t*_*h*+1_). Let us have a look at what happen across epoch (*t*_*h*+1_, *t*_*h*+2_).

The observed punctual event *t*_*h*+1_ can either be,

1. a removed ancestral leaf. Update (3.18) then applies. Subsequently, the number of sampled lineages increases by one and formula (3.17) applies on the next epoch, leading to

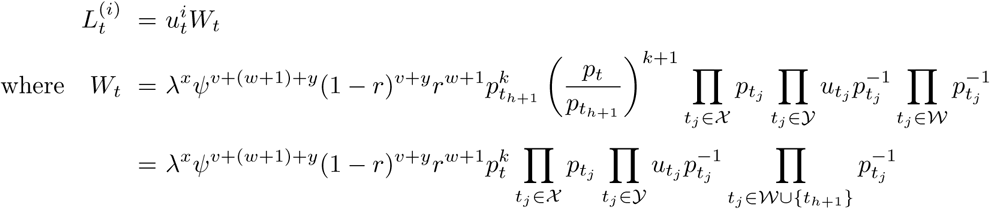
2. a non-removed ancestral leaf. Update (3.19) then applies. Subsequently, the number of sampled lineages increases by one and formula (3.17) applies on the next epoch, leading to

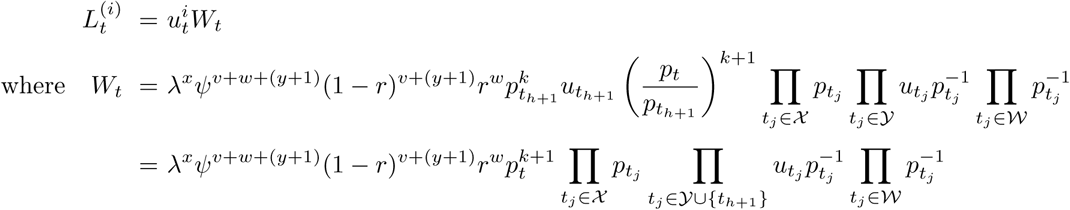
3. a non-removed sampled ancestor along a branch. Update (3.20) then applies. The number of sampled lineages does not changes, and formula (3.17) applies on the next epoch, leading to

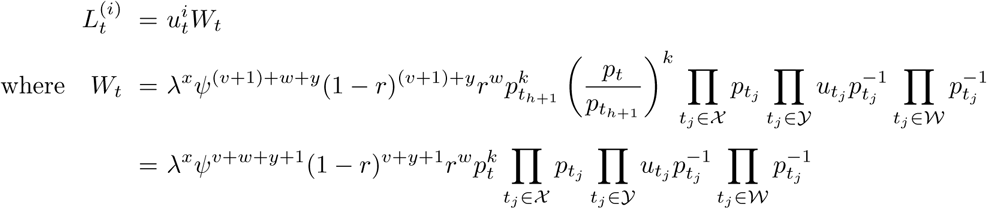
4. a branching event between two sampled lineages. Update (3.21) then applies. The number of sampled lineages decreases by one, and formula (3.17) applies on the next epoch, leading to

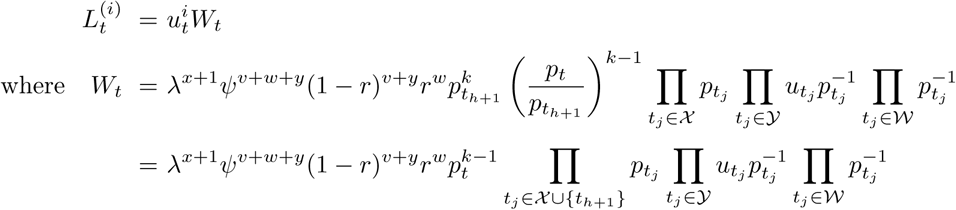

In all four cases, Proposition 3.1 is satisfied across epoch (*t*_*h*+1_, *t*_*h*+2_).

### Appendix D.2. Proof of Proposition 4.3

This Proposition is also proven by induction across the epochs.

We start at *t*_*or*_ = *t*_*n*_ with *k* = 1 lineage. Across epoch (*t*_*n*−1_, *t*_*n*_), applying Proposition 4.1 with *F* (*z*) = 1 and *k* = 1, we get 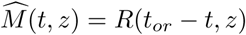, which verifies Proposition 4.3.

Suppose now that Proposition 4.3 is verified across epoch (*t*_*h*_, *t*_*h*+1_) and that we observed, on (*t*_*h*_, *t*_*or*_), *v* sampled ancestors, *w* removed leaves at times *t*_*j*_ ∈ *𝒲, x* branching events at times *t*_*j*_ ∈ *𝒳*, *y* non-removed leaves at times *t*_*j*_ ∈ *𝒴*. Let us have a look at what happens on (*t*_*h*−1_, *t*_*h*_).

Punctual event *t*_*h*_ can either be,

1. a removed leaf. The number of sampled lineages then goes from 1 + *x* − *y* − *w* to *x* − *y* − *w*, and applying update (4.34) followed by Proposition 4.1 leads to

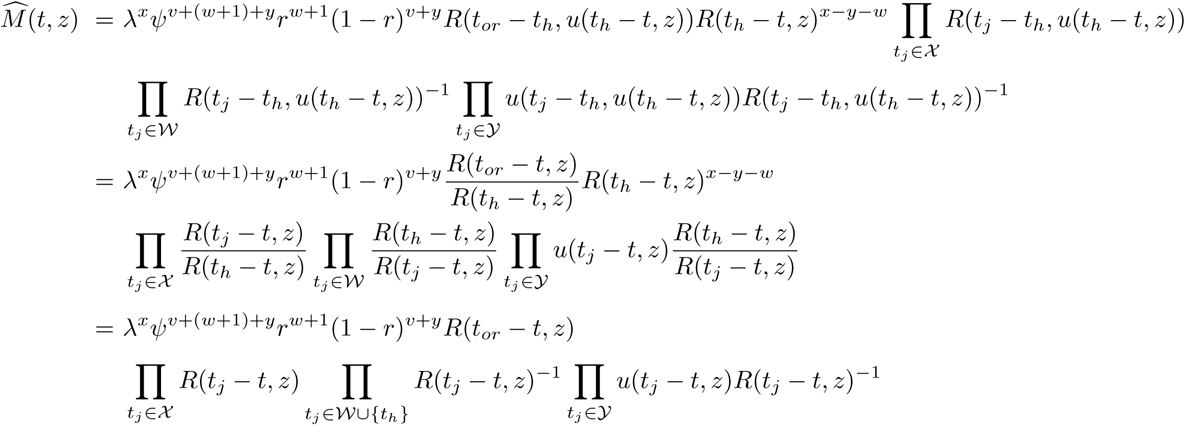

where the first to second equality is detailed in Appendix A, and the second to third comes after canceling out the *R*(*t*_*h*_ − *t, z*).
2. a non-removed leaf. The number of sampled lineages then goes from 1 + *x* − *y* − *w* to *x* − *y* − *w*, and applying update (4.35) followed by Proposition 4.1 leads to

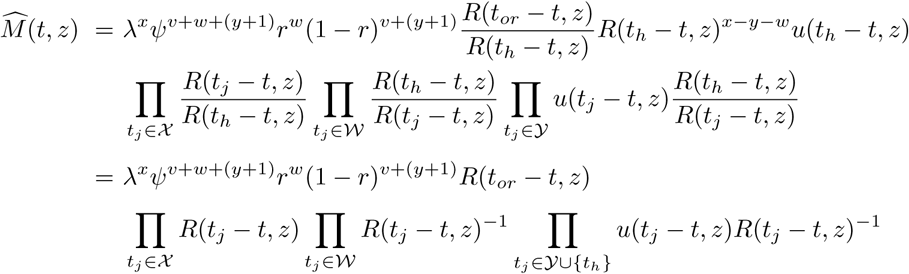
3. a sampled ancestor. The number of sampled lineages then remains unchanged and equal to 1+*x*−*y*−*w*. Applying update (4.36) followed by Proposition 4.1 leads to

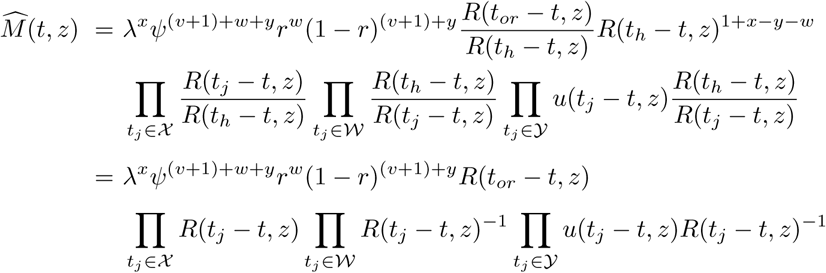
4. a branching time. The number of sampled lineages then goes from 1 + *x* − *y* − *w* to 2 + *x* − *y* − *w*, and applying update (4.39) followed by Proposition 4.1 leads to

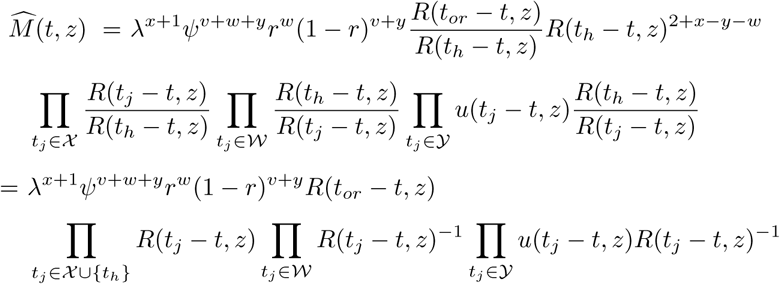

In all these cases, Proposition 4.3 is verified across epoch (*t*_*h*−1_, *t*_*h*_), which ends the proof.

## Appendix E. Using a probability generating function to solve for *L*_*t*_ as well

### Appendix E.1. A slightly different strategy

Recall that *L*_*t*_ verifies the following Master equation,

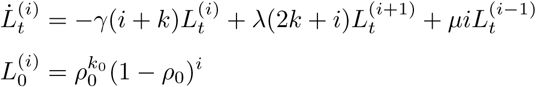

If we introduce the corresponding probability generating function,

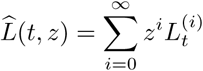

Then the initial condition on *L* translates in 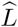 as,

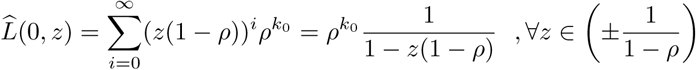

The ODE translates into a PDE, but not as nicely as for *M*_*t*_, see below,

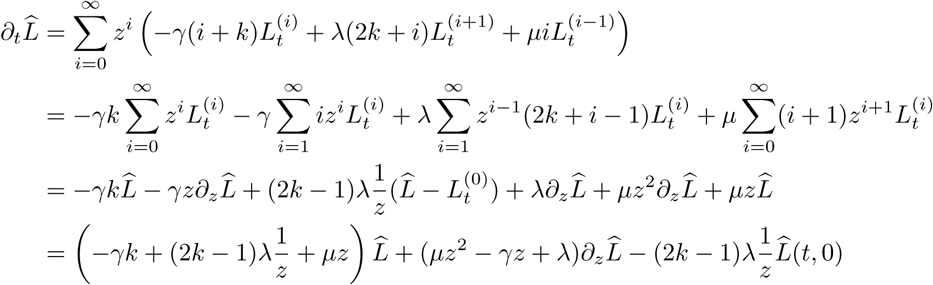

We are thus left with the following PDE problem,

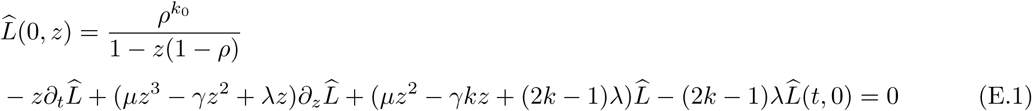

This remaining term with 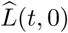 complicates things a little bit. However, the initial condition on 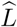 provides us with a first candidate function to satisfy this PDE.

### Appendix E.2. Solution

We introduce below function *f*, and show that it satisfies the PDE problem (E.1).

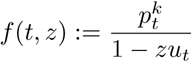

First, we observe that it satisfies the initial condition. We then need to check that it satisfies the PDE, and to do so we expand each of the four components of equation (E.1).

The first one gives us,

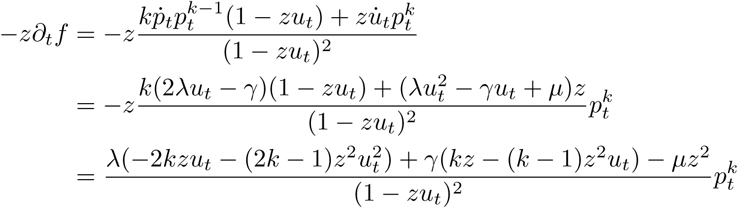

we then turn to the second component,

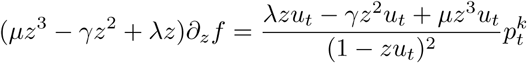

the third one,

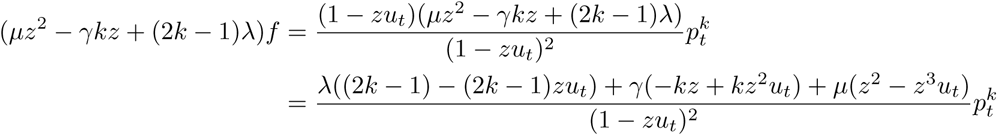

and the fourth and final one,

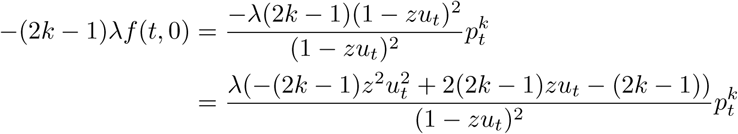

Putting everything together, we can now check that indeed,

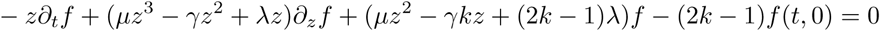

While the branching and *ψ*-sampling with removal updates do not change anything to this solution, all the others do. Further work is thus needed to look for other solutions to this same PDE with different initial conditions.

